# Epigenomic methylome landscape of promoters in vertebrate genomes

**DOI:** 10.64898/2026.03.29.715150

**Authors:** Young Ho Lee, Chul Lee, Erich D. Jarvis, Heebal Kim

## Abstract

Genomic promoters are crucial gene regulatory elements^1,2^. Yet, comparative analyses of promoter architecture have been constrained by the limited resolution of GC-rich regions in short-read-based genome resources^3–6^. The Vertebrate Genomes Project (VGP) provides more complete long-read-based assemblies^7^, which further detect 5-methylcytosine signals directly from PacBio HiFi circular consensus reads^8,9^. Here, we developed a scalable computational framework to characterize DNA methylomes from HiFi data on high-quality Phase I VGP assemblies with RefSeq gene annotations for 82 vertebrate species spanning seven major taxonomic classes: mammals, birds, reptiles, amphibians, lobe-finned fishes, ray-finned fishes, and cartilaginous fishes. We observed a conserved, transcription start site-centered hypomethylation signature in promoters across all vertebrates, and an unexpected hypermethylation signature near gene boundaries that is discordant with transcripts. In addition to this conserved pattern, there were lineage-specific differences in promoter methylation profiles, with birds showing the most diverse patterns. These epigenetic landscapes track phylogenetic relationships more closely than tissue-type methylation differences and infer lineage-dependent widths of core promoters and broader promoters across major vertebrate classes. Our findings establish a comparative epigenomic framework for profiling promoter methylomes from long-read sequencing data.

## Main

Promoters are fundamental regulatory elements that organize transcription initiation^2^ and provide a key substrate for regulatory evolution across vertebrates^1^. Despite decades of work on promoter function and architecture, systematic comparisons across deep vertebrate phylogeny have remained challenging. One reason is technical: promoter regions are often GC-rich with CpG islands, and many older short-read-based genome resources incompletely capture these sequences^3^, complicating analyses that depend on accurate promoter sequence^4–6^. A second reason is practical: large-scale comparative epigenomic surveys have typically been single species focused^10,11^, limiting both phylogenetic breadth and standardization across datasets. As a result, it has been challenging to determine which aspects of promoter organization are broadly conserved across vertebrates and which are lineage-specific.

A conserved promoter hallmark is local hypomethylation around transcription start sites (TSS)^12^, consistent with the idea that reduced CpG methylation makes chromatin accessible to factors for transcription initiation^13^. However, the extent to which promoter-associated methylation patterns differ among major clades, and whether those differences reflect phylogeny rather than sample-specific factors, such as tissue of origin, has remained unclear^14^. In addition, promoter architecture is often discussed in terms of subdomains, including a core promoter that supports transcript initiation and surrounding regions that modulate promoter activity based on transcription factor occupancy^15,16^, chromatin state^17–20^, and sequence context^21,22^. Comparative inference of these features across species has been limited, in part because widely used promoter annotations and functional readouts, such as cap-based TSS mapping^23^ or histone mark profiles^24^, are not uniformly available across vertebrates.

Recent advances in long-read genomics provide an opportunity to revisit promoter evolution at scale. High-quality vertebrate reference assemblies, including those produced by the Vertebrate Genomes Project (VGP), improve access to promoter sequences that were previously fragmented or entirely missing, particularly in GC-rich regions^3,7,25^. In parallel, more recent single-molecule sequencing platforms have been able to simultaneously identify methylation modifications at single-nucleotide resolution by detecting differences in electrochemical profiles. For example, PacBio high-fidelity (HiFi) circular consensus sequencing (CCS) long reads retain kinetic information that directly infers 5-methylcytosine (5mC) at CpG dinucleotides from sequencing data, enabling the construction of DNA methylomes from sequence reads without bisulfite conversion, as used for Illumina short reads^8,26,27^. Together, these advances make it feasible to build standardized, cross-species methylome resources and to compare promoter-associated methylation features across broad phylogenetic sampling.

Here, we infer CpG methylation probabilities (MP) from HiFi reads using a scalable computational framework and integrate these methylomes with curated RefSeq gene annotations for 82 vertebrate species spanning seven major taxonomic classes: mammals, birds, reptiles, amphibians, lobe-finned fishes, ray-finned fishes, and cartilaginous fishes. We observe conserved and lineage-specific promoter methylation profiles that are independent of tissue type. We suggest a quantitative classification of promoter types across species and lineages. Together, our analyses establish a scalable, long-read-derived epigenomic framework for inferring promoter regulatory architecture in assembled vertebrate genomes and provide a comparative view of promoter methylome evolution across vertebrates.

## Results

### Genome-wide human DNA methylome

To define genome-wide methylation patterns across annotated functional contexts, we analyzed PacBio HiFi reads from the near-complete human genome HG002, generated in a sister parallel effort on the human pangenome^28^. Read-level methylation states were inferred from pulse-width and interpulse-duration kinetics. Methylation-called reads were aligned to the telomere-to-telomere human reference T2T-CHM13v2.0 (simply named human sequence 1, hs1)^29^, which provides the RefSeq gene annotations not yet available for the HG002 diploid assemblies. We then used pb-CpG-tools (https://github.com/PacificBiosciences/pb-CpG-tools) to integrate per-read methylation calls across reads mapped to every genomic dinucleotide site where a C is directly followed by a G (CpG) and estimated base-level MP, generating genome-wide human methylation maps (**Fig. 1a; Supplementary Table S1**).

**Figure 1.**
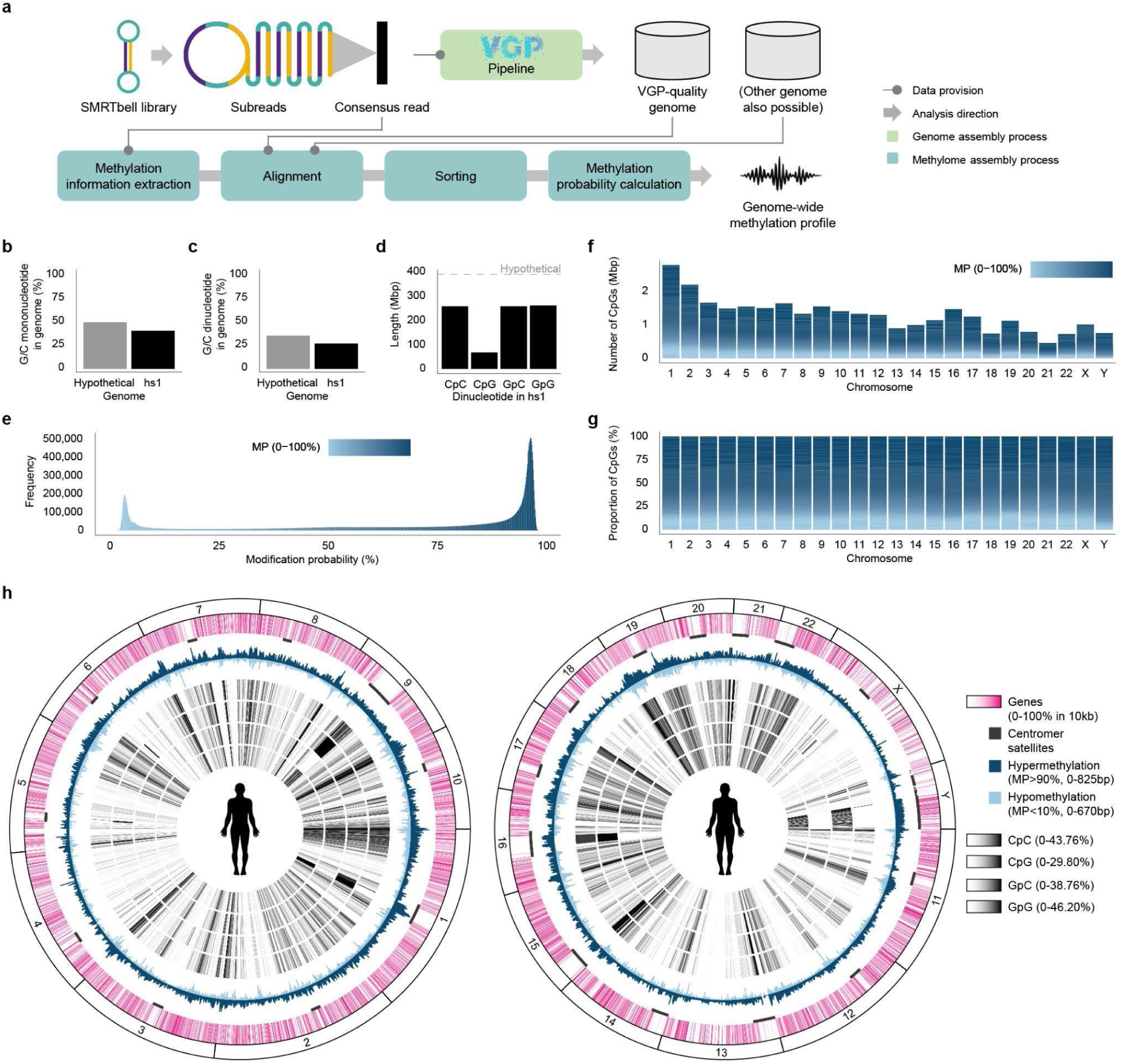
| PacBio HiFi circular consensus reads enable genome-wide inference of the human DNA methylome. **a**, Overview of the methylome inference workflow used in this study. PacBio HiFi circular consensus reads with kinetics tags are aligned to a high-quality reference genome, then sorted and used to calculate methylation probabilities, yielding a genome-wide methylation profile. Arrows indicate the flow of inputs and outputs between genome assembly resources and downstream methylome analyses. **b, c,** Bar plot comparing GC ratio (**a**) and GC-comprised dinucleotide frequency (**b**) in T2T-CHM13v2.0 (hs1) genome with a hypothetical random genome with every base having an equal chance of occurrence. **d,** Bar plot comparing cumulative genomic length occupied by each of the four GC-comprised dinucleotides with indication of the expected occupancy length (10.9375%) in a hypothetical genome with length equal to hs1 in red dashed line for comparison. **e.** Histogram of MP distributions of genomic CpGs. **f, g,** Bar plots of absolute (**f**) and proportionate (**g**) MP distributions of CpGs in each chromosome, with MP value denoted by color. **h**, Circos representation of the T2T-CHM13v2.0 genome (hs1), shown as two panels for readability, larger autosomes 1–10 (left) and smaller autosomes 11–22 plus sex chromosomes X and Y (right). Tracks (outer to inner) display gene annotations, centromeric satellite regions, hypermethylated sites (MP > 0.9), hypomethylated sites (MP < 0.1), and dinucleotide composition for CpC, CpG, GpC, and GpG.

Relative to theoretical expectations under random base composition, hs1 showed lower frequencies of G/C mononucleotides (41% versus 50%) and GC-containing dinucleotides (27.08% versus 37.5%; **Fig. 1b,c**). Among GC-containing dinucleotides (844,275,470 bp; CpC, CpG, GpC, and GpG), CpGs accounted for only 67,792,108 bp (8.03%), far below the 25% expected by chance (**Fig. 1d**). Despite this genome-wide suppression of CpG sites, methylation probabilities at CpGs showed a strongly bimodal distribution across the genome, dominated by hypermethylated sites (**Fig. 1e**). This bimodality was consistent across chromosomes (**Fig. 1f, g**). However, genomic context substantially shaped local methylation patterns: gene-rich regions frequently overlapped CpGs at both hypomethylated (MP < 10%) and hypermethylated (MP > 90%) states, whereas centromeric satellite regions, which were largely gene-poor, were enriched for intermediate methylation states (10 ≤ MP ≤ 90). This marked CpG suppression was also spatially non-random across the genome, with CpGs showing clustered distributions distinct from those of other GC dinucleotides (**Fig. 1h**).

### Genome-wide zebra finch DNA methylome

We next applied the same pipeline to the telomere-to-telomere zebra finch reference bTaeGut7, generated as part of the VGP effort^30^. As an evolutionarily distant comparison to humans and the most complete avian genome assembly available, the zebra finch provides a useful reference for cross-clade methylome analysis. Relative to humans, zebra finches showed sharper intrachromosomal transitions between hypermethylated and hypomethylated regions, with gene-rich microchromosomes tending to be more hypermethylated (**Extended Data Fig. 1**). In contrast, like the human Y chromosome, the gene-poor zebra finch W chromosome was also enriched for hypermethylated regions. Like the human X chromosome, the zebra finch Y chromosome was relatively hypomethylated compared with the autosomes (**Extended Data Fig. 1**; **Fig. 1h**). Overall, localized intervals of elevated gene density or hypomethylation were interspersed with GC-rich sequence contexts.

### DNA methylation of regulatory elements

To obtain a genome-wide view of methylation across regulatory elements, we profiled CpG methylation within ±10 kb of centers of annotated elements in GRCh38/hg38 at single-base resolution. We used hg38 rather than hs1 for this analysis because ENCODE candidate *cis*-regulatory element annotations are currently only available for hg38. Regulatory coordinates were compiled from RefSeq and ENCODE annotations for promoters, enhancers, and silencers. Across all categories (promoters, enhancers, and silencers), both mean and median MP profiles showed a pronounced V-shaped dip centered on each element, extending for approximately 2.5 kb on either side (**Fig. 2a,b**). To assess the significance of this dip-like pattern, we compared MP distributions at each base position within the ±10 kb window with the genome-wide MP distribution using total variation distance (d_TV_), a non-parametric approach appropriate for the strongly non-normal genome-wide MP distribution (**Fig. 1e**). This analysis confirmed that the central decrease in MP was statistically significant across regulatory element classes, indicating that local hypomethylation is a pervasive epigenomic feature of human regulatory elements.

**Figure 2.**
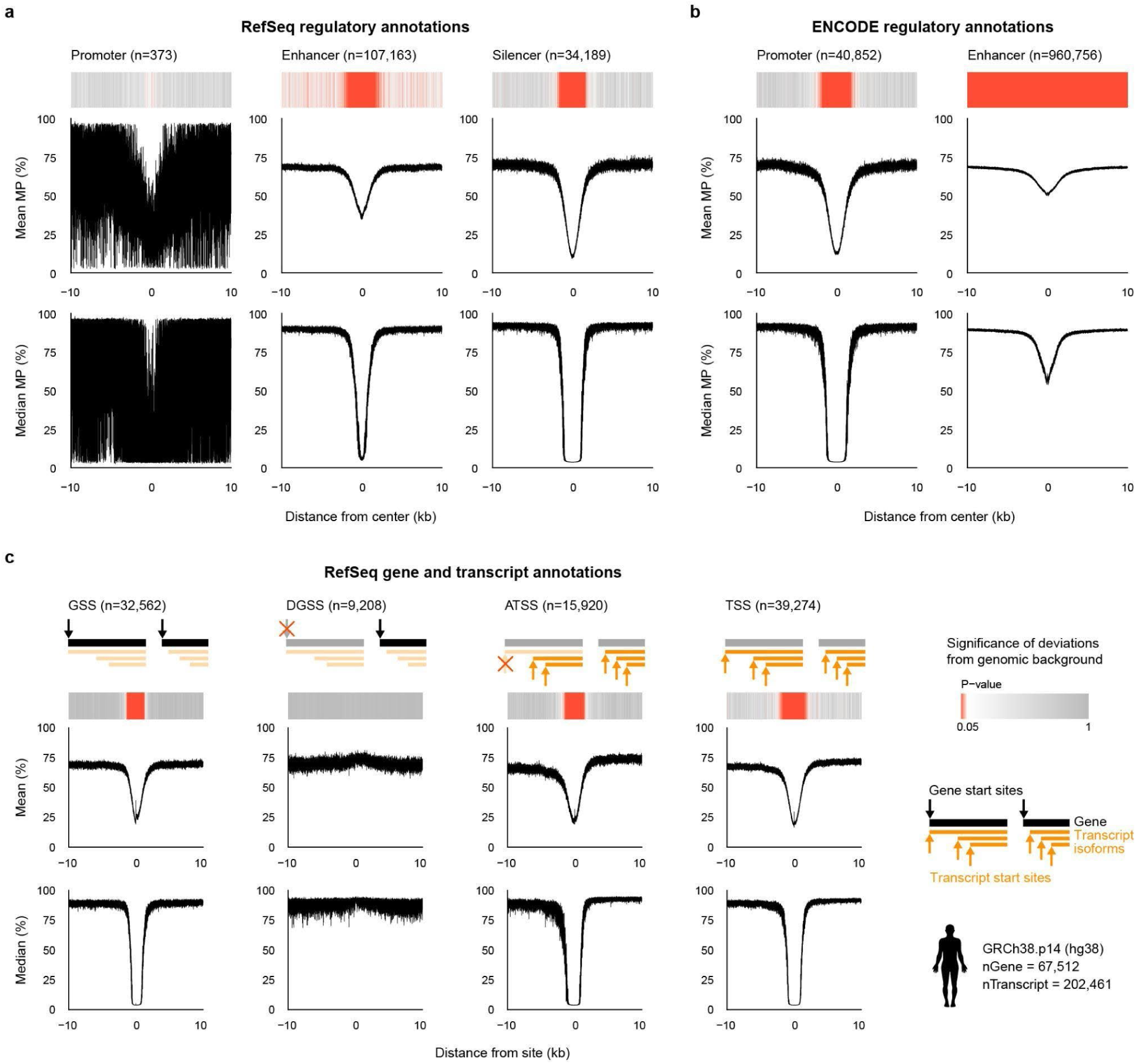
| Diverse human regulatory annotations converge on a TSS-centered hypomethylation signature. **a**, Line plots show mean (top) and median (bottom) MP of CpGs as a function of distance from the center for RefSeq regulator (promoter, enhancer, and silencer) annotation centers, **b**, ENCODE candidate promoter and enhancer centers, and **c,** RefSeq gene-related start sites, namely, gene start sites (GSS), discordant gene start sites (DGSS), alternative transcript start sites (ATSS), and transcript start sites (TSS). CpG MP profiles are inferred from PacBio HiFi kinetics and mapped to the human reference genome (GRCh38.p14), summarized around annotated regulatory elements and start sites of genes and transcripts within a ±10 kb window. Heatmaps show distance-resolved significance of deviations from a genome-wide background, quantified by d_TV_ and converted to p-values (Methods).

We next examined whether the methylation signature near genes changed. We generated MP profiles centered on gene start sites (GSS) where the RefSeq annotation and transcript start sites align, discordant gene start sites (DGSS) where the RefSeq annotation start is upstream of the first transcript start, alternative transcript start sites (ATSS), and all transcript start sites (TSS) (**Fig. 2c**; **Supplementary Table S2**). GSS, ATSS, and TSS showed a significant V-shaped decrease in MP centered on the annotated start site, closely resembling the pattern observed across annotated regulatory elements, whereas the DGSS did not. These results support the accuracy of RefSeq start-site annotations, except for the likely discordant annotated gene start sites, and further localize promoter-associated hypomethylation within the gene 5’ boundaries. By contrast, analogous analyses at termination sites, including gene termination site (GTS), discordant gene termination sites (DGTS), alternative transcript termination sites (ATTS), and transcript termination sites (TTS), revealed only an abrupt local decrease at the boundary itself, without the extended region of significant hypomethylation seen at start sites (**Extended Data Fig. 2**; **Supplementary Table S2**). Similar boundary-associated methylation patterns were also observed in the human genome hs1 using its corresponding RefSeq gene annotations (**Extended Data Fig. 3**; **Supplementary Table S2**).

To examine promoter methylation at single-gene resolution, we analyzed CpG methylation profiles within ±10 kb of the GSS and TSS of two ubiquitously expressed human housekeeping genes, *ACTB* and *GAPDH*. Both loci showed near-complete hypomethylation around the TSS of these genes and for the surrounding genes, with almost all CpGs in the local promoter region remaining below 25% in MP (**Extended Data Figs. 6** and **7a,b**).

### Vertebrate promoter methylome landscapes

We also asked whether these boundary-associated methylation patterns were conserved in an evolutionarily distant vertebrate. We therefore profiled methylation around gene and transcript boundaries in two zebra finch assemblies, bTaeGut1.4.pri and bTaeGut7.mat, using their respective RefSeq annotations (**Extended Data Figs. 4,5**; **Supplementary Table S2**). In both assemblies, 5’ gene and transcript boundaries retained the pronounced V-shaped dip seen in human start-site profiles, indicating that promoter-associated hypomethylation is conserved across these vertebrate lineages. However, the DGSS profile in bTaeGut7.mat displayed periodic fluctuations that produced regularly spaced spikes and also influenced the aggregate GSS profile (**Extended Data Fig. 5**). To minimize this assembly-specific noise in downstream analyses, we used TSS-centered methylation profiles.

To investigate evolutionary variation in promoter methylation across vertebrates, we extended our methylome assembly and TSS-centered MP profiling framework to 86 additional vertebrate genomes from VGP Phase I, together with the two human and two zebra finch genomes analyzed above, yielding an initial set of 88 species. This dataset spanned all seven major vertebrate classes (**Supplementary Table S1**). Seven genomes showing artefactual MP profiles were excluded from downstream analyses, leaving 83 genomes, including 34 mammals, 16 birds, 13 ray-finned fishes, 9 reptiles, 6 amphibians, 4 cartilaginous fishes, and 1 lobe-finned fish for comparative profiling.

Across species, mean promoter MP profiles consistently retained the characteristic dip observed in humans, but the shape of this feature varied across vertebrate classes. Birds showed the shallowest decrease, followed by mammals and reptiles, whereas amphibians and fishes exhibited steeper flanks and shorter troughs, despite substantial within-class variation, particularly among amphibians and reptiles (**Fig. 3**). Baseline MP levels flanking the dip also differed by class: birds, reptiles, and fishes showed lower and more gradual declines, amphibians showed a distinctly steeper 5’-side descent, and mammals showed the most symmetrical profiles. These class-level differences were even more apparent in median profiles, consistent with promoter MP divergence being a genome-wide property rather than one driven by a small subset of loci.

**Figure 3.**
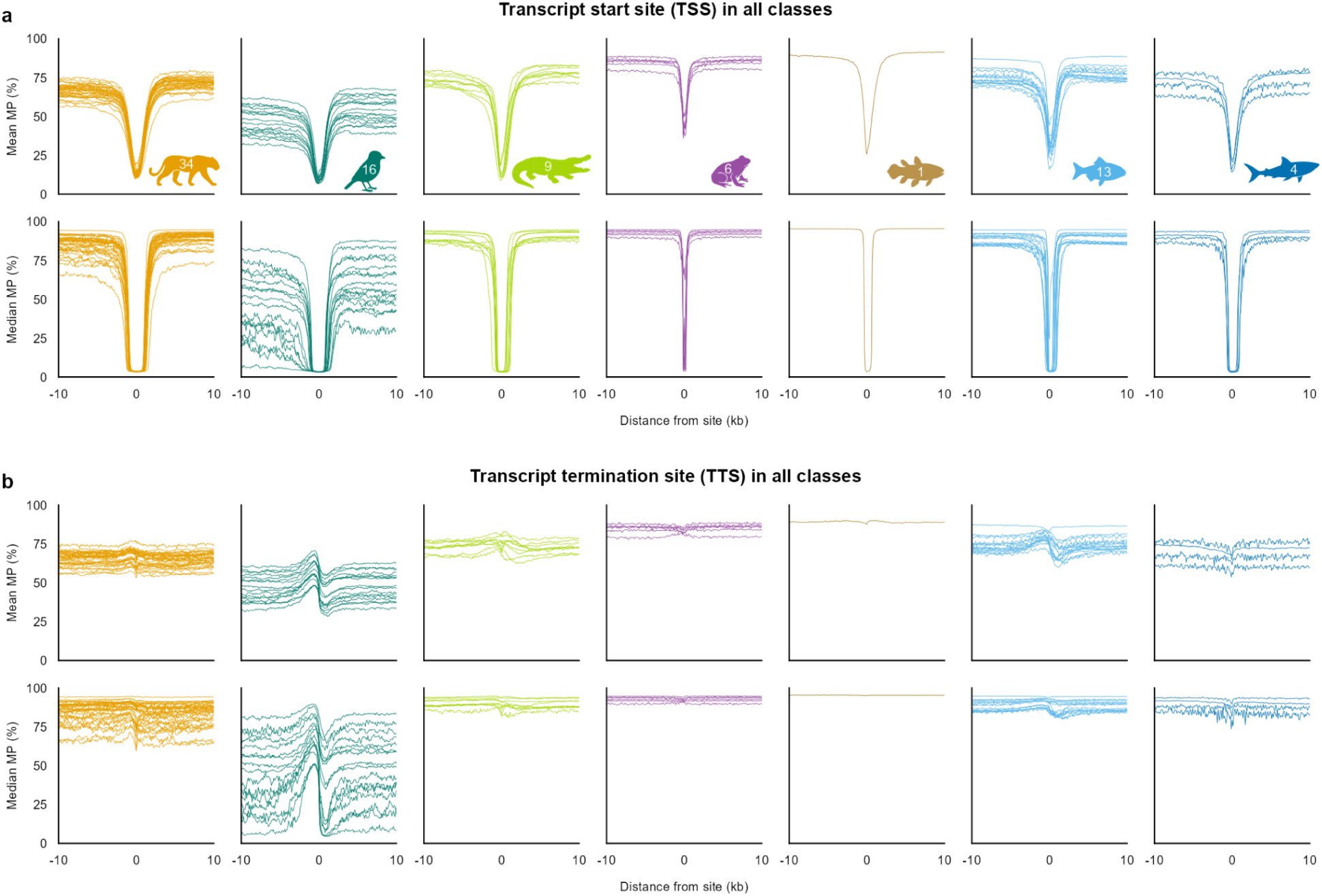
| TSS and TTS methylation profiles are conserved across vertebrates but differ by class. **a,b**, Mean and median methylation probability (MP) of CpGs as a function of distance from transcription start sites (TSS; **a**) and transcription termination sites (TTS; **b**) in VGP Phase I genomes. Each curve represents one genome, grouped by major vertebrate class (left to right): all classes (# species = 88, # genome = 90), mammals (35, 36), birds (18, 19), reptiles (9, 9), amphibians (6, 6), lobe-finned fishes (1, 1), ray-finned fishes (16, 16), and cartilaginous fishes (4, 4). Across classes, TSS-centered hypomethylation is preserved, while profile shape and breadth vary among lineages.

Similar to what was done on humans, we examined orthologous *ACTB* and *GAPDH* genes across vertebrate species spanning multiple classes. Promoter-associated hypomethylation was broadly preserved, although the span and local structure of low-MP regions varied substantially among species, with some loci showing broader hypomethylated tracts or additional low-MP islands (**Extended Data Fig. 7c,d**). Because these comparisons are based on only two genes, however, they are insufficient to infer class-level differences and may instead reflect locus-specific effects of neighboring functional elements.

### Lineage-specific stronger than tissue-specific

To quantify class-level divergence in promoter methylation, we performed Uniform Manifold Approximation and Projection (UMAP)^31,32^ on MP profiles across vertebrate genomes. Using the mean CpG methylation probability within ±10 kb of the TSS, the first two UMAP components separated genomes largely by vertebrate class (**Fig. 4a**). TTS-centered profiles showed weaker class separation, but combining TSS and TTS profiles produced a more distinct clustering pattern by class. UMAP profiles of the median and SD showed results similar to those of the mean, but were less prevalent (**Extended Data Fig. 8a,c**).

**Figure 4.**
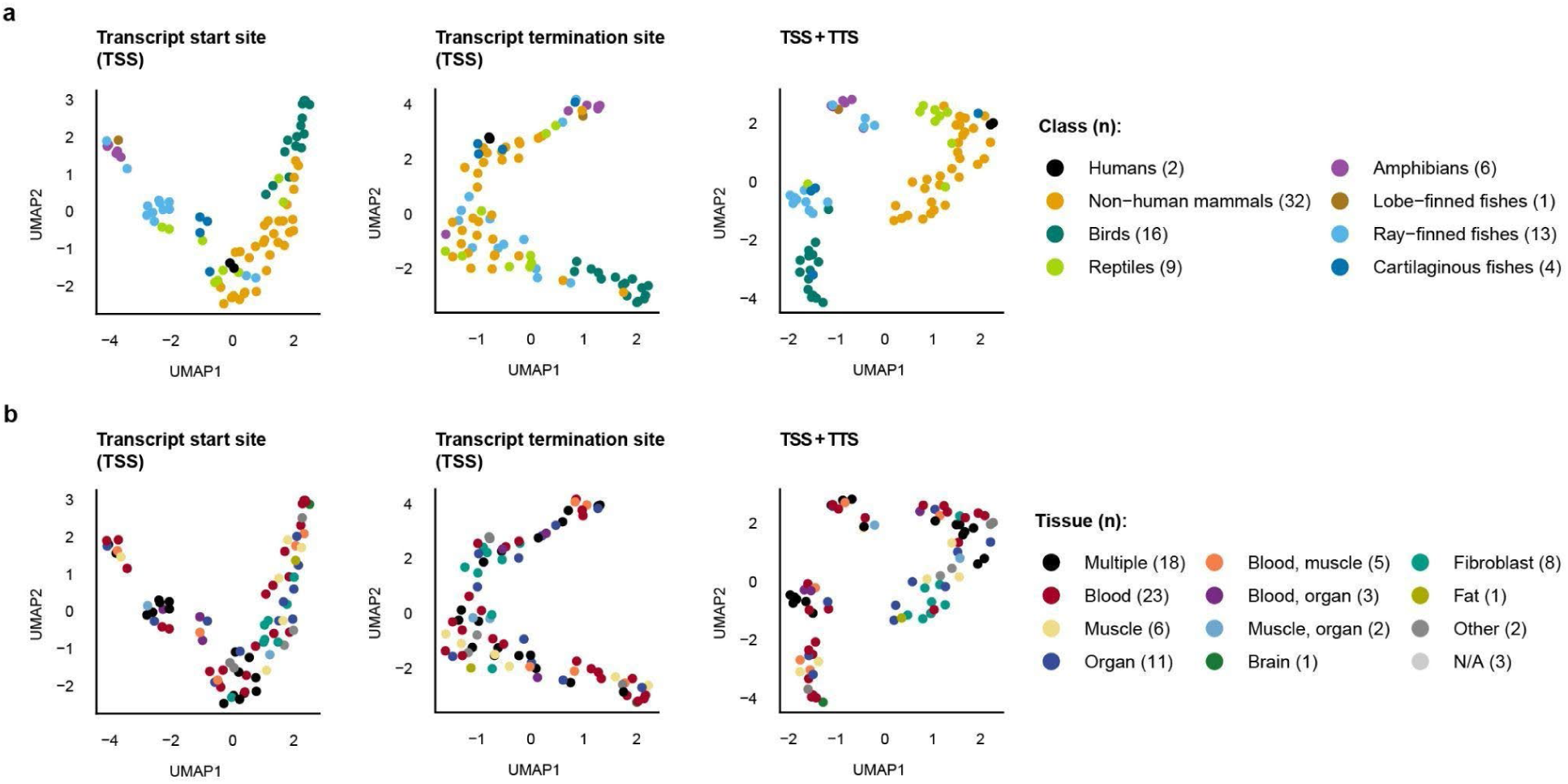
| Promoter-associated methylation clusters by lineage, more than tissue of origin. **a**, UMAP analysis of mean MP profiles summarized within ±10 kb of annotated transcription start sites (TSS), transcription termination sites (TTS), or both combined (TSS+TTS) across 90 VGP genomes. Each point represents one genome, colored by taxonomic class using the same color codes as in Fig. 3. **b,** The same UMAP projections shown in **a** are colored by tissue of origin. Across feature sets and genomic contexts, samples separate primarily by lineage rather than tissue.

We observed segregation in varying degrees among a few sub-lineages for which we had sufficient species data. For example, although the primate order showed subclustering profiles indistinct from the rest of the mammal class at both TSS and TTS sites (**Extended Data Fig. 9**), passeriform birds (i.e., songbirds) displayed a subclustering profile from the rest of the mammals, explained by their lower baseline MP levels (**Extended Data Fig. 10**).

Because VGP samples were collected and sequenced independently for each species (**Supplementary Table 1**), we considered whether tissue-of-origin differences might contribute to the observed clustering. When genomes in the UMAP were labelled by tissue type source, only modest tissue-associated structure was apparent (**Fig. 4b, Extended Data Fig. 8b,d**). The most consistent grouping involved fibroblast-derived samples, which were predominantly used for mammals, and a weaker tendency was observed among species represented by mixed-tissue or whole-organism samples, mainly in fishes and small amphibians. Given these sampling asymmetries, tissue-related effects cannot be excluded entirely. Overall, the observed tissue signal was much weaker than the separation by vertebrate class, suggesting that lineage identity is a stronger organizing axis of promoter methylome variation than tissue type in this dataset.

### Broad– and core-promoters across species

To obtain a higher resolution profile of methylation around the TSS, we analyzed the mean MP profile revealed a localized mini W-shaped feature centered on the TSS across species, consisting of a narrow central rise flanked by two minima that together spanned approximately 100 bp in human (**Fig. 5a**). This feature was asymmetric, with a deeper and steeper valley on the 5’ side, suggesting that methylation asymmetry around the TSS may mark the core promoter, defined as a region of nearly 0 methylation and inferred as active chromatin (**Fig. 5c**). To test this possibility, we compared mean MP on the 5’ and 3’ sides of the TSS in 10-bp windows. We assessed positional significance using the corresponding CpG MP distributions. The 5’ side remained significantly less methylated than the 3’ side across an interval consistent with a 150-bp core promoter in human, defining an epigenetically asymmetric region at the promoter center (**Fig. 5d**). To determine whether this epigenetic boundary framework extended beyond humans, we applied the same general promoter breadth and core-promoter estimation strategy to the zebra finch. Although the asymmetry was less visually prominent than in humans (**Fig. 5b**), the 5’ and 3’ sides of the TSS remained significantly different, defining a 200-bp core promoter in zebra finch (**Fig. 5e**).

**Figure 5.**
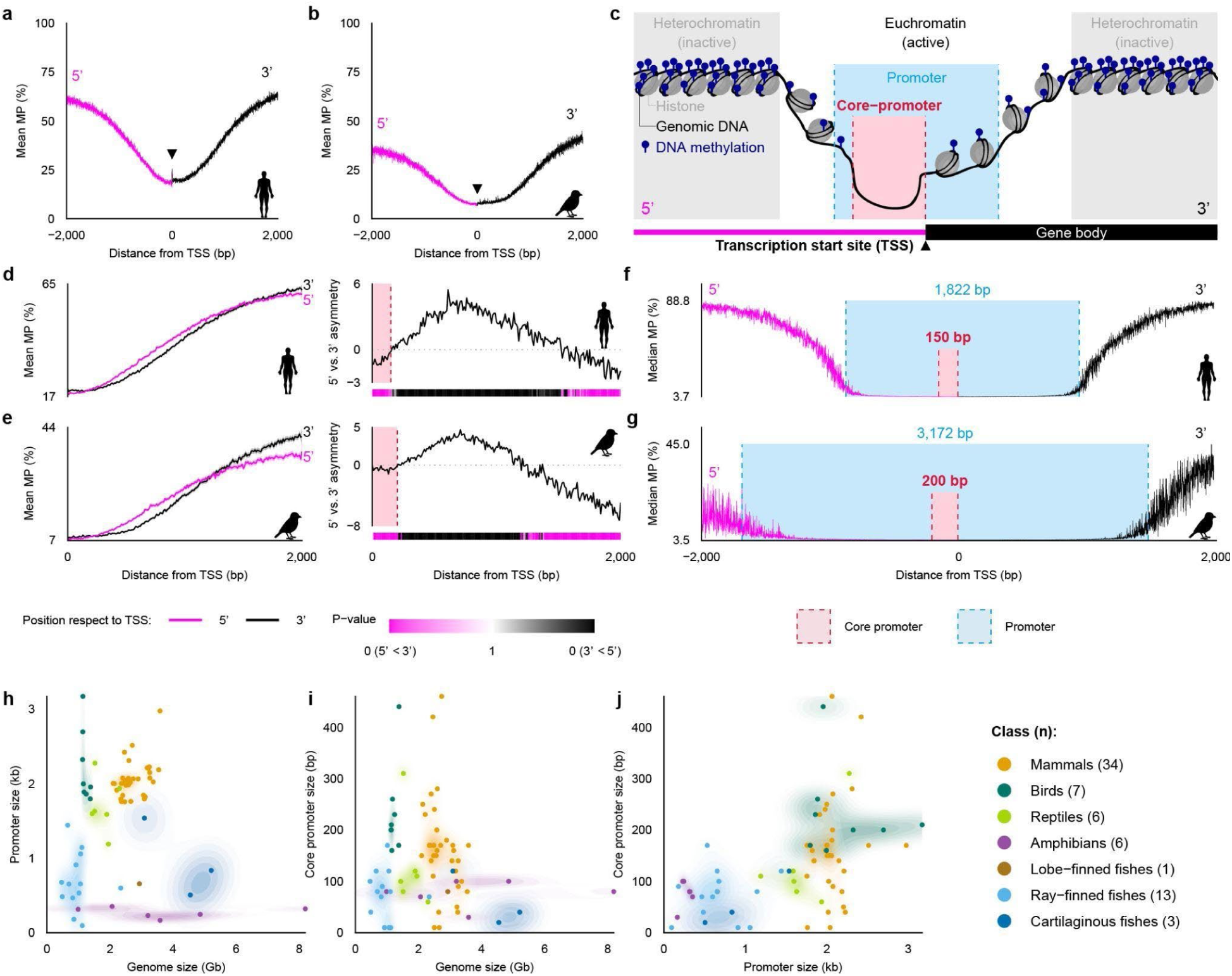
| TSS-centered 5′–3′ methylation asymmetry and a local methylation dip delineate promoter boundaries. **a,b**, Mean methylation probability (MP) of CpGs as a function of distance from RefSeq transcription start sites (TSS) within a ±2 kb window for human (T2T-CHM13v2.0, **a**) and zebra finch (bTaeGut7.mat, **b**). Profiles are shown separately for the 5’ (magenta) and 3’ (black) sides of the TSS, and the arrowhead marks the TSS position. **c**, Schematic illustrating a promoter region around a TSS, highlighting the methylation-defined promoter (blue) and core promoter (red) in the context of chromatin accessibility. **d,e**, Left, side-specific mean MP profiles plotted against absolute distance from the TSS for human (**d**) and zebra finch (**e**). Right, 5’–3’ asymmetry computed from the difference between the side-specific MP profiles, with a distance-resolved heatmap indicating the significance of asymmetry in 10-bp windows based on a total variation distance test between 5’ and 3’ MP distributions (Methods); the color scale encodes both p-value and direction (5’<3’ versus 3’<5’). **f,g**, Median MP profiles within a ±2 kb window, with methylation-defined promoter boundaries (blue dashed lines and shading) and core promoter boundaries (red dashed box) estimated as described in Methods, shown for human (**f**) and zebra finch (**g**). Promoter size and core promoter size were defined from the methylation dip and 5’–3’ asymmetry, respectively (Methods). **h-j,** Scatterplot showing relationships between methylation-defined promoter size versus genome size (**h**), methylation-defined core promoter size versus genome size (**i**), and methylation-defined promoter size versus core promoter size (**j**) across species, with points colored by taxonomic class.

We next estimated the broader promoter region from the median TSS methylation profile, which showed a steeper and more sharply bounded decline in MP than the mean profile. We identified two boundary positions flanking the TSS that maximized the contrast in per-base median MP between the putative promoter and surrounding non-promoter sequence within a ±10 kb interval. This analysis estimated a general promoter breadth of 1,822 bp in the human genome (**Fig. 5f**) and a substantially wider promoter estimate of 3,172 bp in the zebra finch genome (**Fig. 5g**).

### Promoter breadth of vertebrate classes

The differences in general and core promoters between humans and zebra finches led us to consider whether there are overall lineage-specific differences. We applied this framework across all VGP species in our dataset to estimate general promoter and core-promoter breadths for each genome. These estimates revealed broad class-level differences, with mammals, birds, and reptiles generally showing larger promoter regions than amphibians, ray-finned fishes, and cartilaginous fishes (**Fig. 5h-j, 6**; **Supplementary Table S1**). Birds showed the broadest promoter and core-promoter intervals, whereas amphibians showed the narrowest promoter breadth. Across classes, the overall 5’-3’ asymmetry around the TSS was preserved (**Fig. 6**).

**Figure 6.**
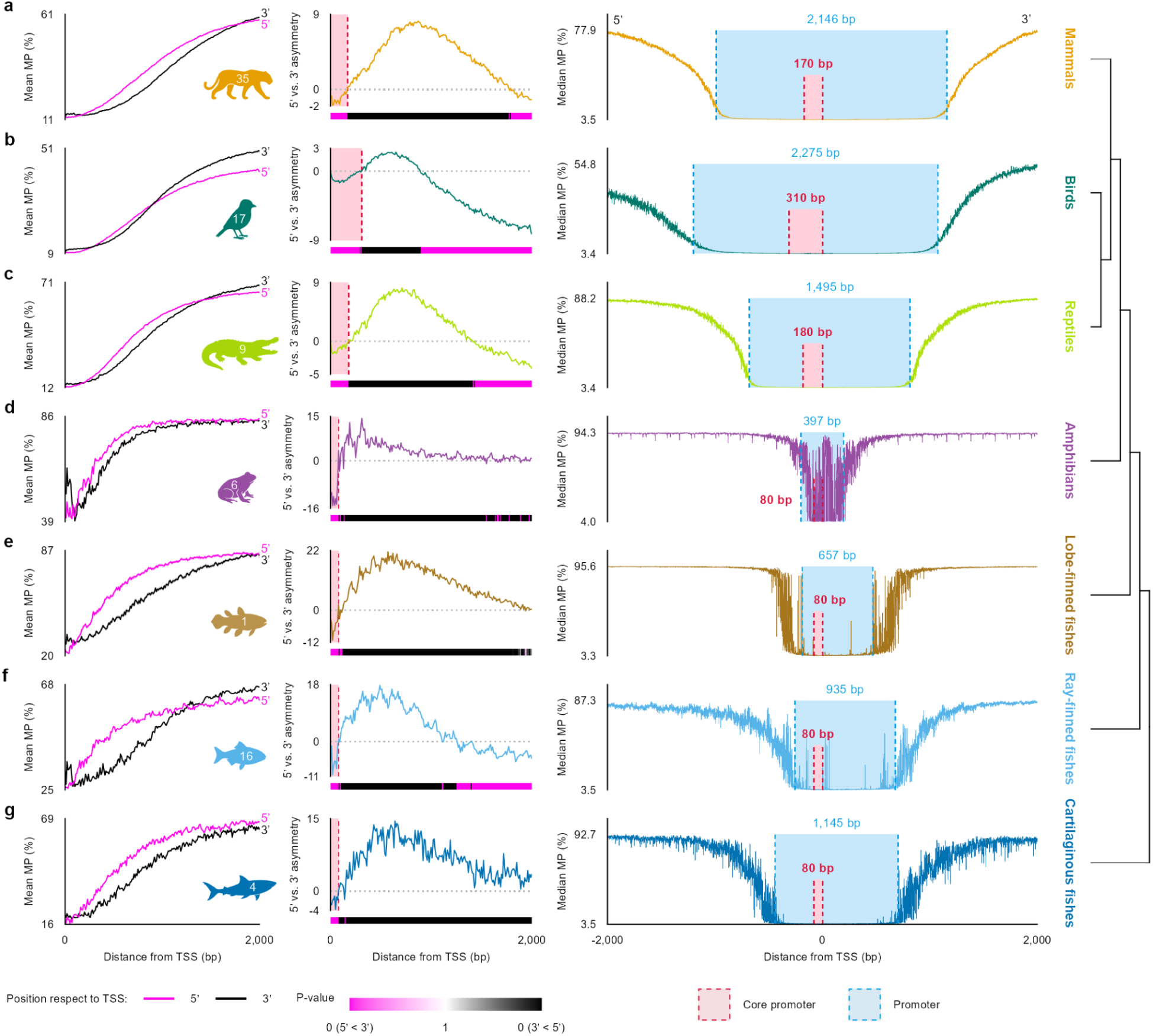
| Core promoter extent and promoter breadth vary phylogenetically across vertebrate classes. **a–g**, Class-level summaries of methylation probability (MP) around transcription start sites (TSS). Left, mean MP as a function of distance from the TSS, shown separately for the 5’ and 3’ sides. Middle, 5’–3’ asymmetry computed from side-specific MP differences, with a heatmap showing windowed significance (10-bp windows) based on a total variation distance test comparing 5’ and 3’ MP distributions. Right, median MP profiles with estimated broader promoter and core promoter boundaries indicated (values shown in panels). Major vertebrate classes are shown as mammals (**a**), birds (**b**), reptiles (**c**), amphibians (**d**), lobe-finned fishes (**e**), ray-finned fishes (**f**), and cartilaginous fishes (**g**) using the same color codes as in Fig. 3.

Promoter breadth did not scale with genome size, but species class did cluster according to shared promoter breadth and genome size (**Fig. 5h-j**). For example, mammals consistently showed larger promoter regions than amphibians despite similar genome sizes, and birds showed larger promoter and core-promoter breadths than fishes despite their similarly compact genomes. General promoter and core promoter breadths were only weakly correlated overall, with birds tending to exceed mammals in both measures (**Fig. 5j**). Together, these results indicate that promoter architecture captures phylogenetically structured epigenetic differences that are not explained simply by genome size, consistent with the class-level separation observed in our methylation clustering analyses.

General promoter breadth varied more strongly among classes than core promoter breadth, although the two measures were weakly correlated overall (**Fig. 6**). Reptiles occupied an intermediate position, whereas amphibians, lobe-finned fish, ray-finned fish, and cartilaginous fish generally showed shorter promoter and core-promoter intervals than mammals and birds. These class-level profiles did not indicate strong within-class smoothing in all groups, suggesting substantial heterogeneity among member species. Even so, the aggregate promoter and core-promoter breadths, which capture lineage-level trends (**Fig. 6**), are consistent with the species-level patterns observed (**Fig. 5h-j**).

## Discussion

Promoter-proximal methylation profiles, inferred from PacBio HiFi kinetics and centered on annotated transcription start sites, are conserved across 82 vertebrate species representing the breadth of the vertebrate family tree, with lineage-specific architectures across different vertebrate classes. Leveraging fine-scale 5’–3’ methylation asymmetry and local undermethylation near TSS, we propose methylation-derived, operational estimates of core-like promoter extent and methylation-defined general promoter breadth across vertebrate genomes.

The substantial depletion of methylation at CpGs is consistent with prior observations across vertebrates^33^. Such depletion likely reflects evolutionary forces beyond GC-specific pressures alone, and may include processes related to methylation-associated mutability. In CpG-rich regions, hypomethylation has been reported^34,35^, thus elevated CpG density may itself contribute to chromatin accessibility and dosage regulation through gradations in methylation. CpG density has also been linked to promoter-associated properties^36^, including associations between promoter CpG levels and lifespan in vertebrates^37^. Although we observed that hypomethylated sites frequently overlap genes, gene presence alone does not dictate methylation state, as methylation can vary across orthologous loci^38^, amplicons^35^, and repeats^39^.

When aggregated genome-wide, a methylation feature manifests as a broad V-shaped depression in the mean MP profile around promoter sites, consistent with the notion that hypomethylation accompanies chromatin states permissive for transcription initiation^40^. Mirroring the methylation V-shaped dips, future tests will be necessary to assess possible correlations with open chromatin peaks at the GSS, DGSS, ATSS, and TSS. We also need to consider multimodal integrations of transcriptome and epigenome patterns to identify gene and transcript boundaries beyond homologous-sequence-based searches in gene annotation.

Similar depressions are observed at other regulatory elements, supporting overall agreement between methylation landscapes and functional element positions identified by NCBI RefSeq and ENCODE. The promoter-associated drop appears less pronounced for RefSeq promoter annotations, which may reflect the limited number of candidate promoters. One possible interpretation of lineage-specific differences in the drop in promoter methylation is that trough duration serves as an operational proxy for methylation-defined promoter breadth.

We also find that the degree of within-lineage concordance varies across clades, with some lineages clustering more tightly than others. In certain cases, species segregate down to the order level based on promoter MP features, including baseline levels, for example, among passerines. In contrast, other orders, such as primates, are comparatively indistinguishable from other members of the class in these promoter-level MP summaries, despite well-established phenotypic and genomic divergence^41–43^. One possible interpretation is that epigenomic and genomic evolutionary rates can be partially decoupled, and that lineage-associated methylation architectures may be shaped by constraints that are not strictly mirrored by sequence divergence alone. Mechanistically, such patterns could be consistent with models involving targeted demethylation outside DNA replication, often referred to as active demethylation^44,45^. More generally, because methylation can influence transcription factor occupancy^46,47^ and local mutational processes^48,49^, it could contribute to regulatory divergence, warranting focused mechanistic follow-up.

At higher resolution near the TSS, the subtle W-shaped methylation structure is reminiscent of classical promoter architecture^50^. Given the bimodal MP distribution, these patterns suggest that CpG methylation proportions differ across promoter subregions, potentially supporting more graded, modulatory behavior rather than a strictly binary model. While some studies also implicitly support this non-binary model by showing that transcriptional machinery and DNA-binding factors can engage promoter regions across diverse methylation states^38^.

The 5’–3’ asymmetry embedded in the W-shaped profile, characterized by uneven valleys on the upstream and downstream sides, provides an operational signal to define a core-like promoter extent uniformly. One hypothesis we propose is that this asymmetry reflects a transition in epigenetic state across the initiation boundary, consistent with broader links between gene-body methylation and expression regulation reported in prior literature^51^. Importantly, the general promoter and core-like extents we infer are methylation-defined operational estimates and are not, by definition, equivalent to transcription initiation breadth as measured by cap-based TSS mapping in human and mouse^52,53^. Establishing how methylation-defined promoter breadth relates to canonical promoter classes will require direct benchmarking against orthogonal functional readouts, including CAGE or related TSS mapping assays, as well as chromatin signatures such as ATAC-seq and H3K4me3, in species where such data can be generated or compiled.

Using this epigenetically defined framework, we can provide general estimates of promoter and core promoter sizes computed solely from methylation data. These estimates provide a comparative reference for how promoter-associated methylation architecture varies across major vertebrate classes and species. Promoter architecture properties, including methylation, have been shown to vary with gene function and context^54^, including cell-^55^, tissue-^56^, or stage-specific expression^57,58^, and species-lineage, indicating that different species evolved genome-wide regulatory properties. The lack of correlation between the promoter methylation profile and genome size indicates that methylation-defined promoter architecture does not scale simply with overall genome expansion.

Birds show comparatively broad methylation-defined promoter regions in our estimates, despite their compact genome sizes. One possible explanation is that avian genome organization, including gene-dense microchromosomes and elevated recombination rates, could shape promoter sequence composition and regulatory landscapes, which in turn influence methylation architecture. Testing this hypothesis will require analyses stratified by microchromosomes versus macrochromosomes and matched regulatory datasets across tissues. Consistent with this possibility, prior studies have reported heightened epigenetic plasticity and variable CpG properties across subsets of bird promoters, particularly for stress– or immune-related genes^59^, suggesting that broad promoter-associated methylation architectures may be driven by specific functional categories rather than by uniform methylation across all genes.

Our study has some limitations. Although we ruled out tissue source thus far, species samples differ in developmental stage and tissue state (cell culture vs tissue), and these factors can covary with lineage due to sampling practices in VGP. TSS positions rely on RefSeq annotations whose completeness and precision vary across species, potentially affecting fine-scale asymmetry and boundary estimates. Although this is not a problem with the ENCODE annotations. Our fixed ±10 kb window may truncate long-range promoter-associated patterns in some taxa, whereas expanding it would introduce additional regulatory contexts. These considerations motivate follow-up analyses using matched tissues or cell types within selected lineages, and benchmarking methylation-derived promoter metrics against orthogonal functional assays where available.

Nonetheless, by constructing and analyzing high-resolution methylomes across 82 VGP species, this study provides one of the most extensive standardized comparative views of vertebrate promoter methylation to date, and profiles many species epigenetically for the first time. Using a uniform analytical framework, we identify lineage-associated promoter methylation signatures and derive methylation-based, operational estimates of promoter and core-like promoter extents across major vertebrate classes. Because these methylomes are inferred directly from publicly available genome assembly reads provided by VGP collaborators, our approach reduces experimental barriers and provides a scalable path to extend comparative epigenomic analyses to additional species. In conclusion, this resource and set of quantitative metrics provide a cross-vertebrate reference for testing hypotheses about the evolution of promoter architecture, improving promoter annotation in more complete genomes, and enabling systematic comparative studies that were previously impractical at this phylogenetic scale.

## Extended Data Figures

**Extended Data Fig. 1.**
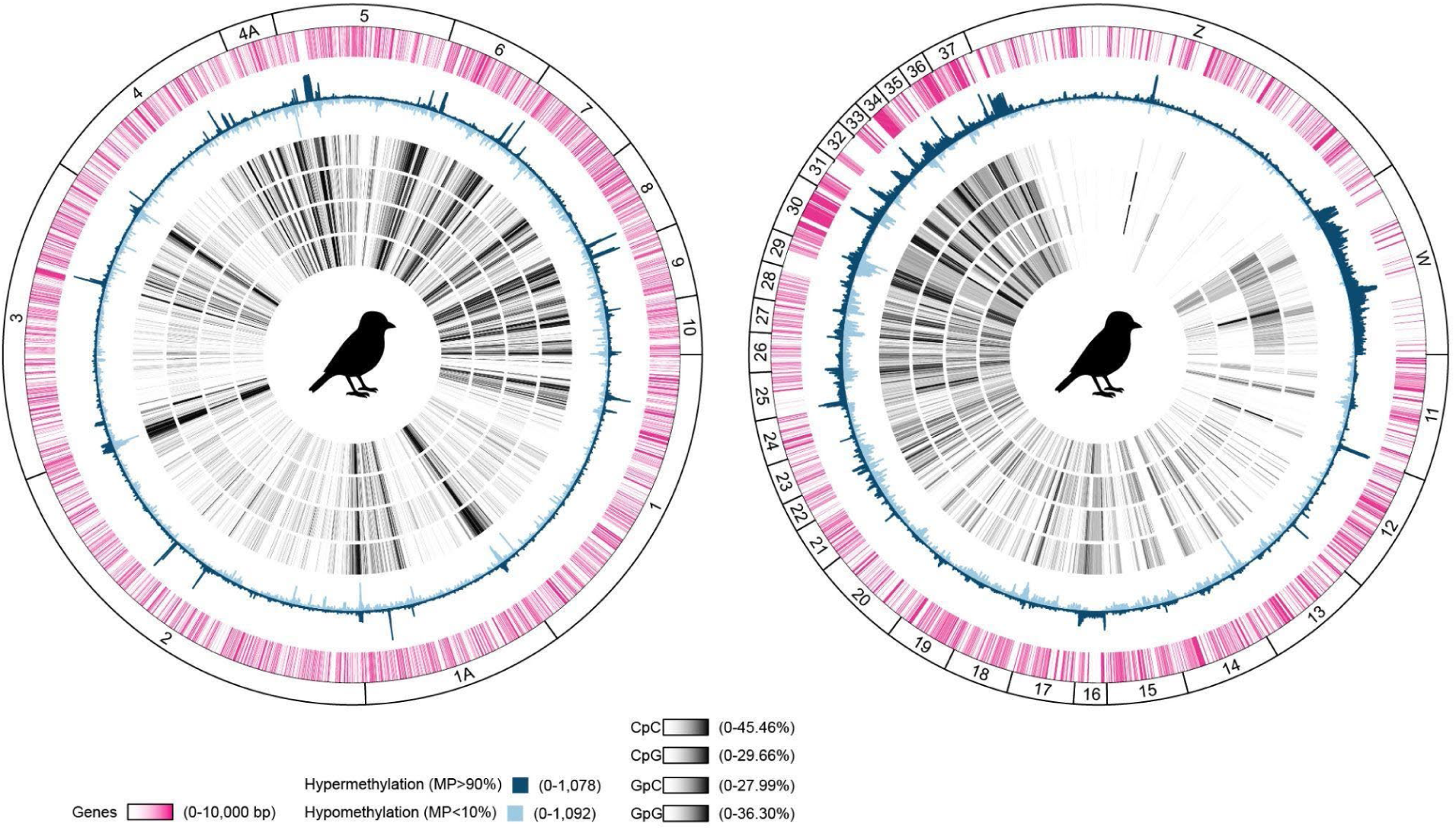
| Zebra finch DNA methylome. Circos plots about the zebra finch genome bTaeGut7.mat separated into upper (chromosomes 1-10, 1A, 4A) (left) and lower (chromosomes 11-37, Z/W) (right) halves with each track representing (from outermost) genes, hyper– (MP>90%) and hypomethylated sites (MP>10%), and GC-comprised dinucleotides (CpC, CpG, GpC, and GpG).

**Extended Data Fig. 2.**
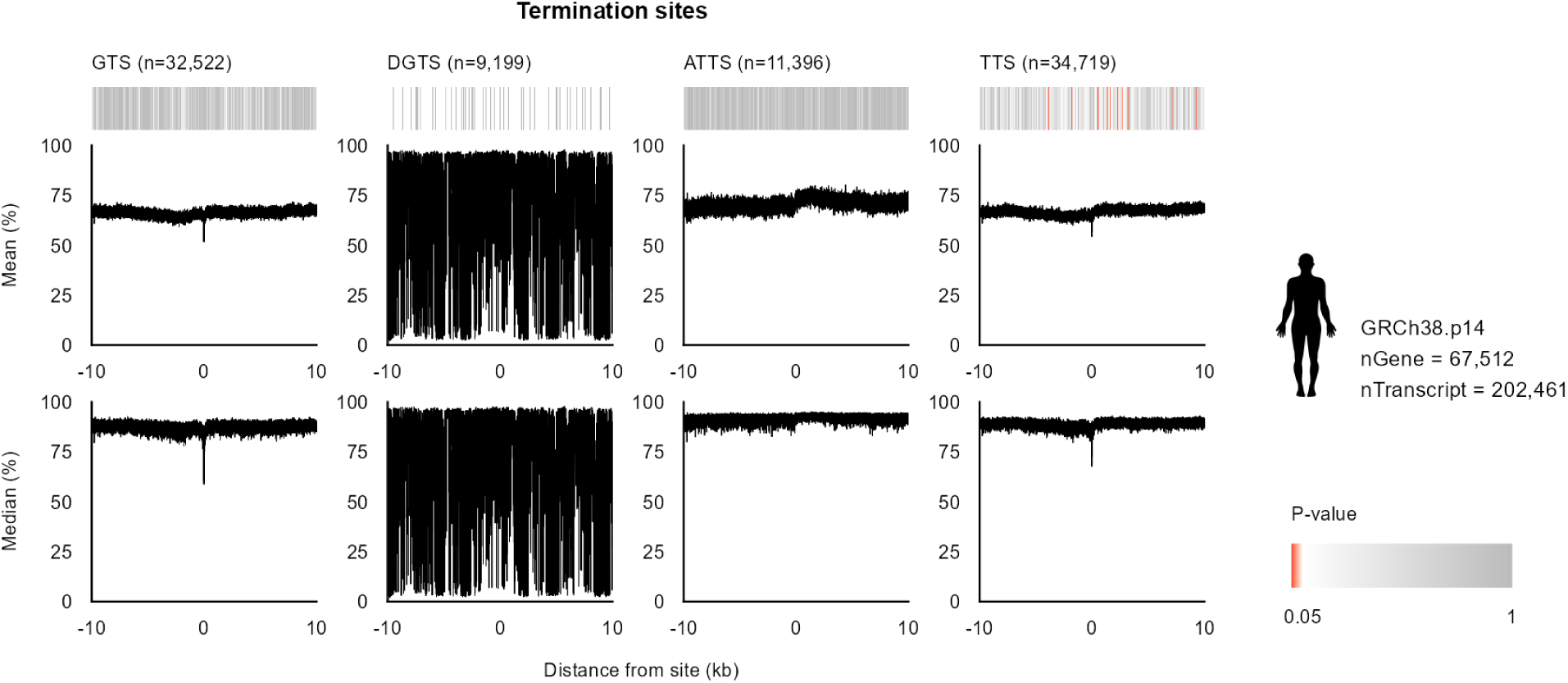
| Various significant methylation properties are displayed at start and end sites of estimated transcriptions in the human reference genome hg38. Deviation of methylation probability (MP) profiles from the genome-wide background around putative regulatory centers in the traditional human reference genome hg38. Heatmaps summarize the significance of deviations quantified by d_TV_ and converted to *p-*values; darker red indicates smaller *p*-values. Line plots show the mean and median MP of CpGs as a function of distance from start and terminal sites of genes (GSS, GTS), discordant start and terminal sites of genes compared to that of transcripts (DGSS), start and terminal sites of transcripts (TSS, TTS), and start and terminal sites of alternative transcripts (ATSS, ATTS) across a ±10 kb window.

**Extended Data Fig. 3.**
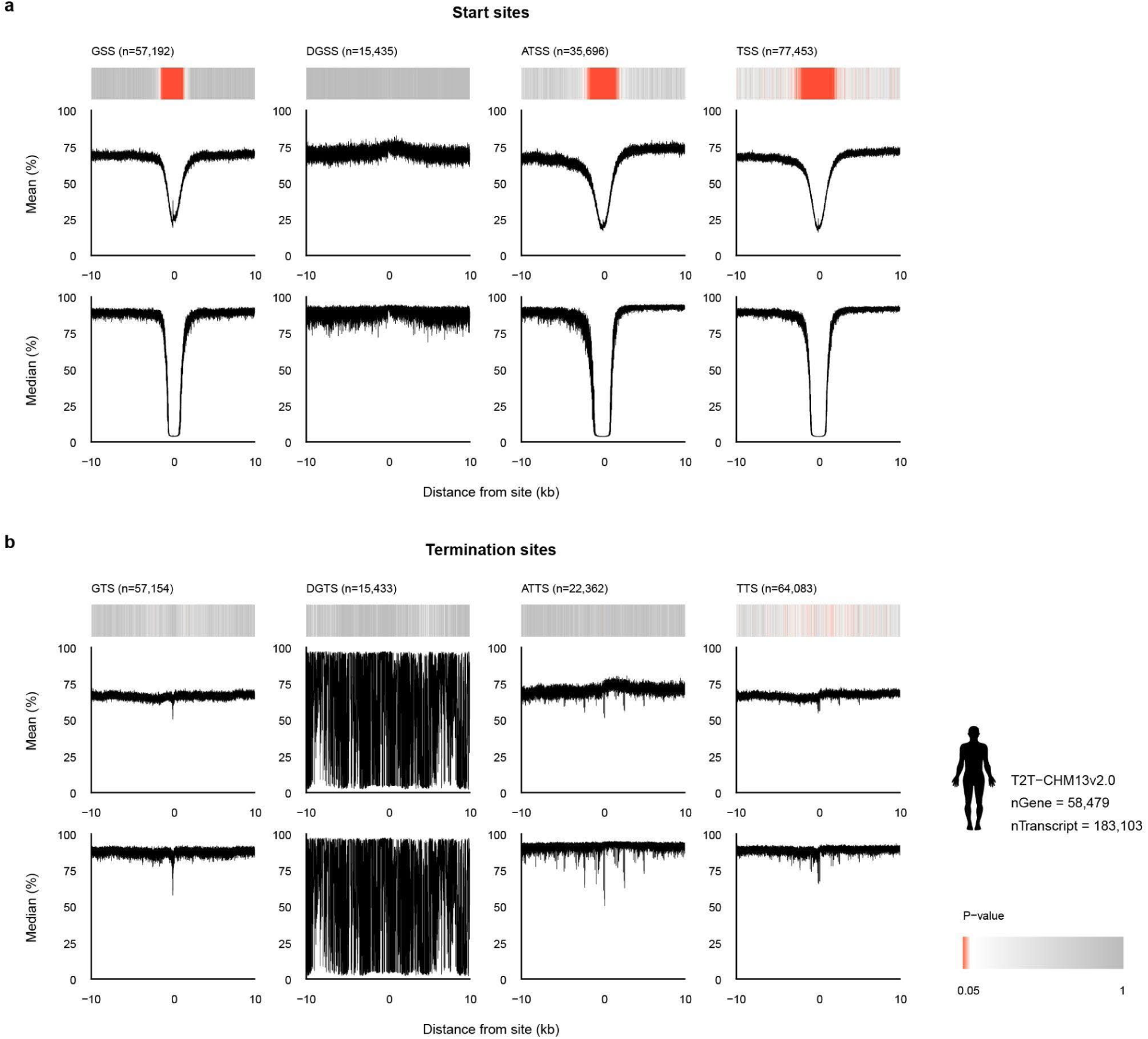
| Human promoter methylation patterns are replicated across promoter profiles of the human RefSeq gene annotation of the telomere-to-telomere genome (hs1). **a**, Methylation probabilities of gene and transcript start sites. **b,** Methylation probabilities of gene and transcript termination sites. Identical format as **Extended Data** Fig. 3, for the human telomere-to-telomere assembly hs1.

**Extended Data Fig. 4.**
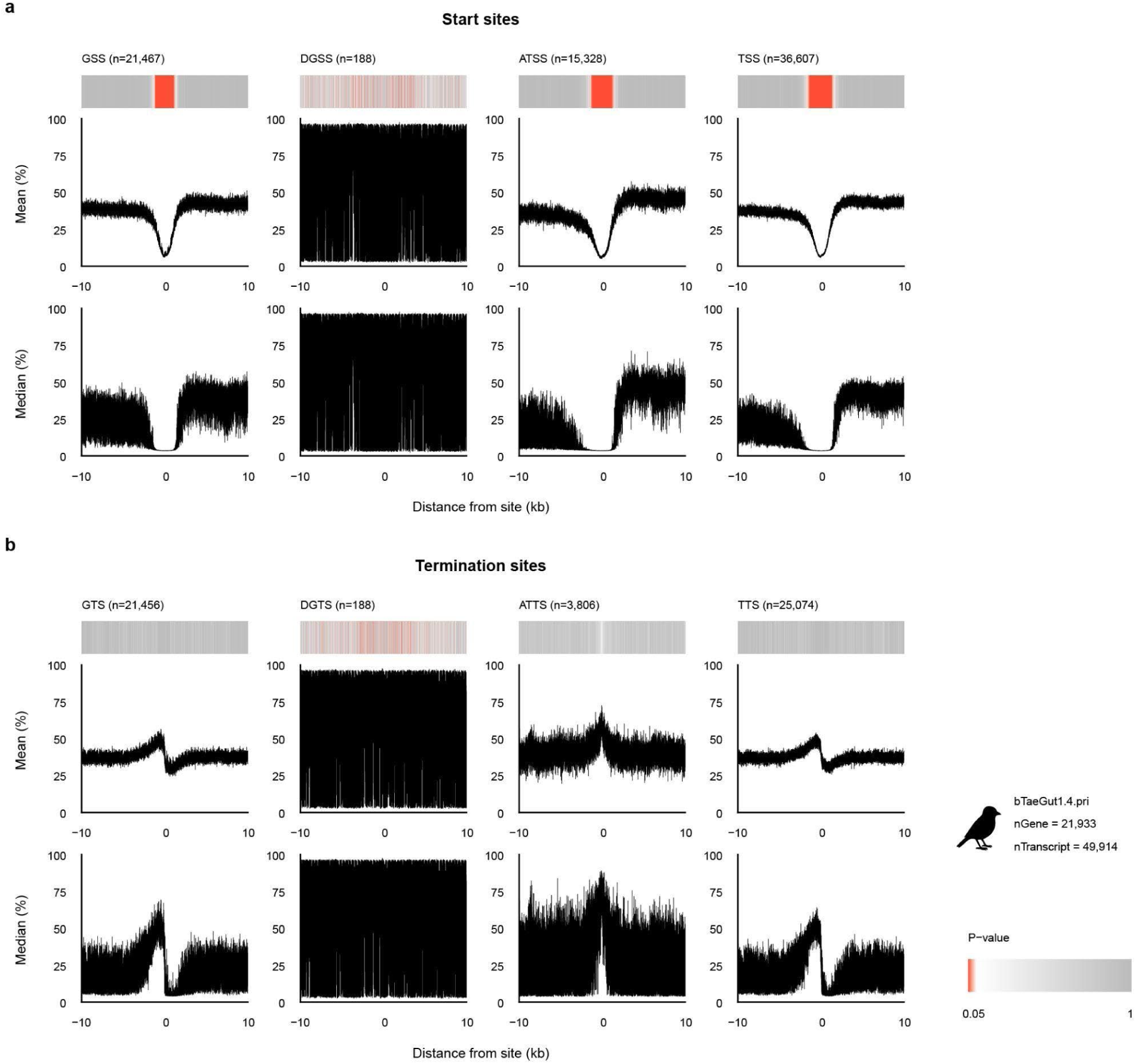
| A different methylation pattern ensues in promoters of a non-human genome – zebra finch (bTaeGut1.4.pri). **a**, Methylation probabilities of gene and transcript start sites. **b,** Methylation probabilities of gene and transcript termination sites. Identical format as **Extended Data** Fig. 3, for the previous zebra finch reference genome bTaeGut1.4.pri.

**Extended Data Fig. 5.**
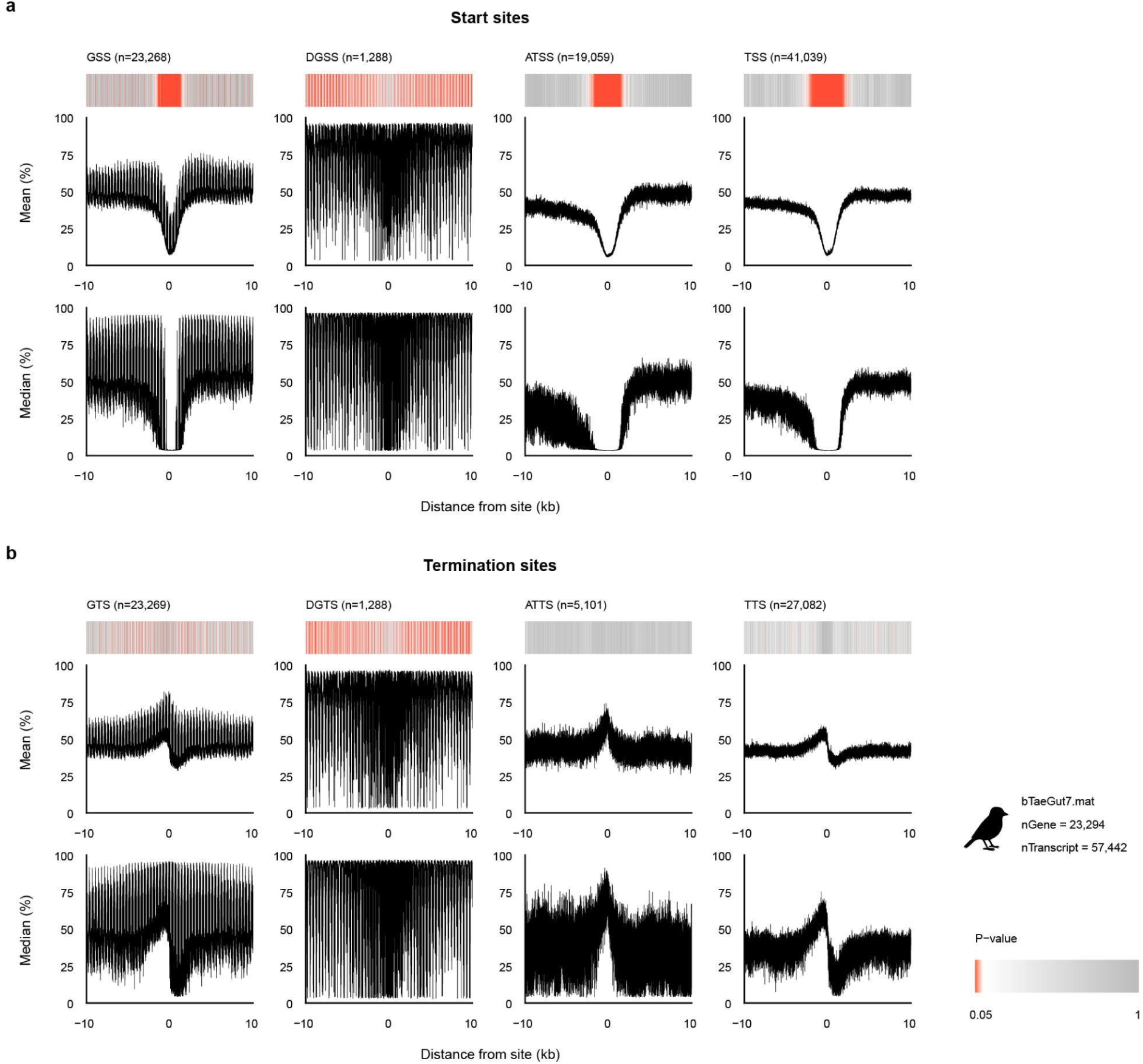
| Unusual aberrations are found in the telomere-to-telomere genome of zebra finch (bTaeGut7.mat) arising from its unique gene start sites. **a**, Methylation probabilities of gene and transcript start sites. **b,** Methylation probabilities of gene and transcript termination sites. Identical format as **Extended Data** Fig. 3, for the zebra finch telomere-to-telomere assembly bTaeGut7.mat.

**Extended Data Fig. 6.**
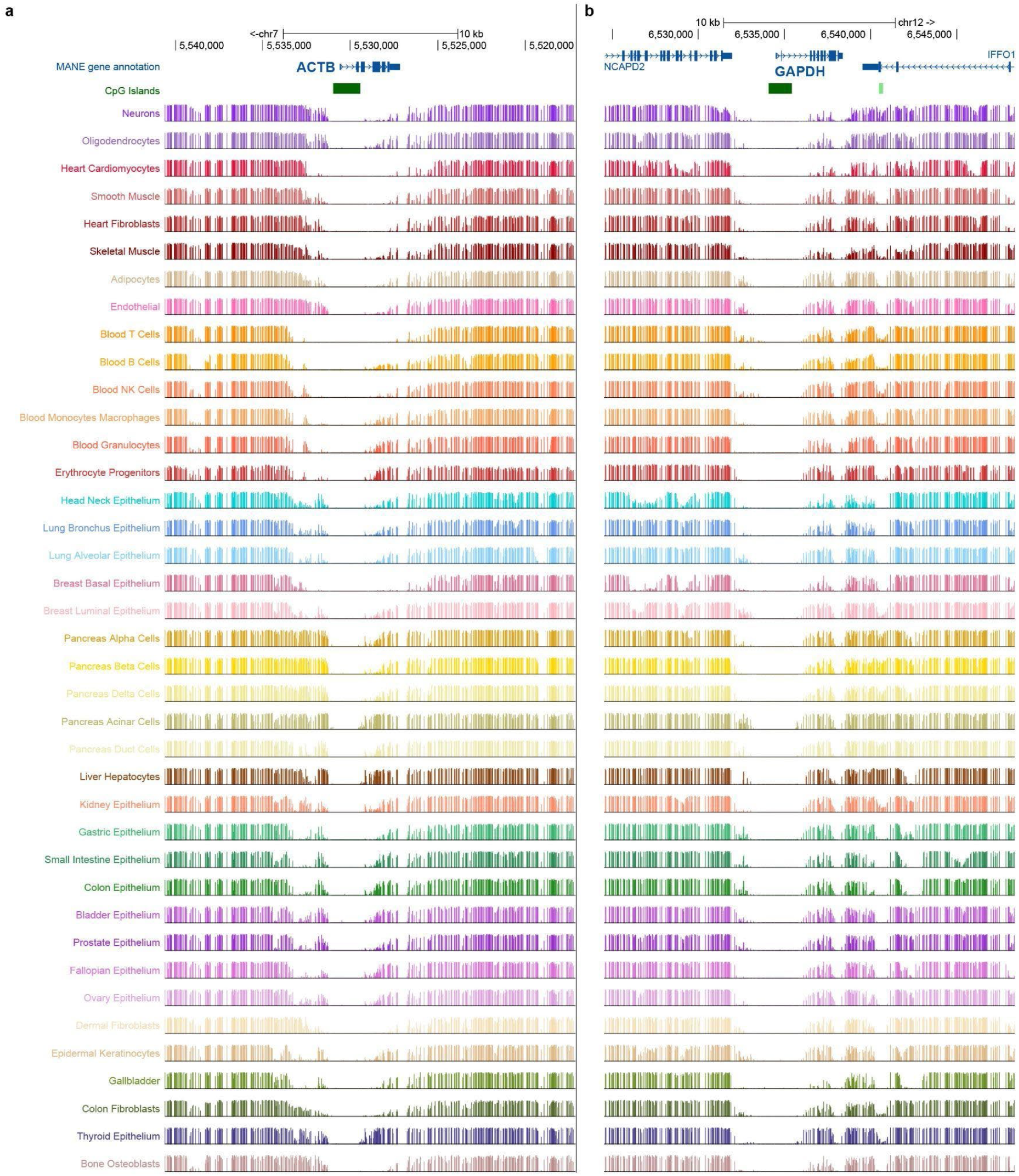
| Methylation profile at constitutively expressed genes. **a, b**, MP levels of whole genome bisulfite sequencing data from various tissues on the hg38 genome with respect to physical distances from TSS for genes *ACTB* (**a**) and *GAPDH* (**b**) within ±10,000 bp intervals of the site. Methylation data visualized from the UCSC genome browser^60–62^.

**Extended Data Fig. 7.**
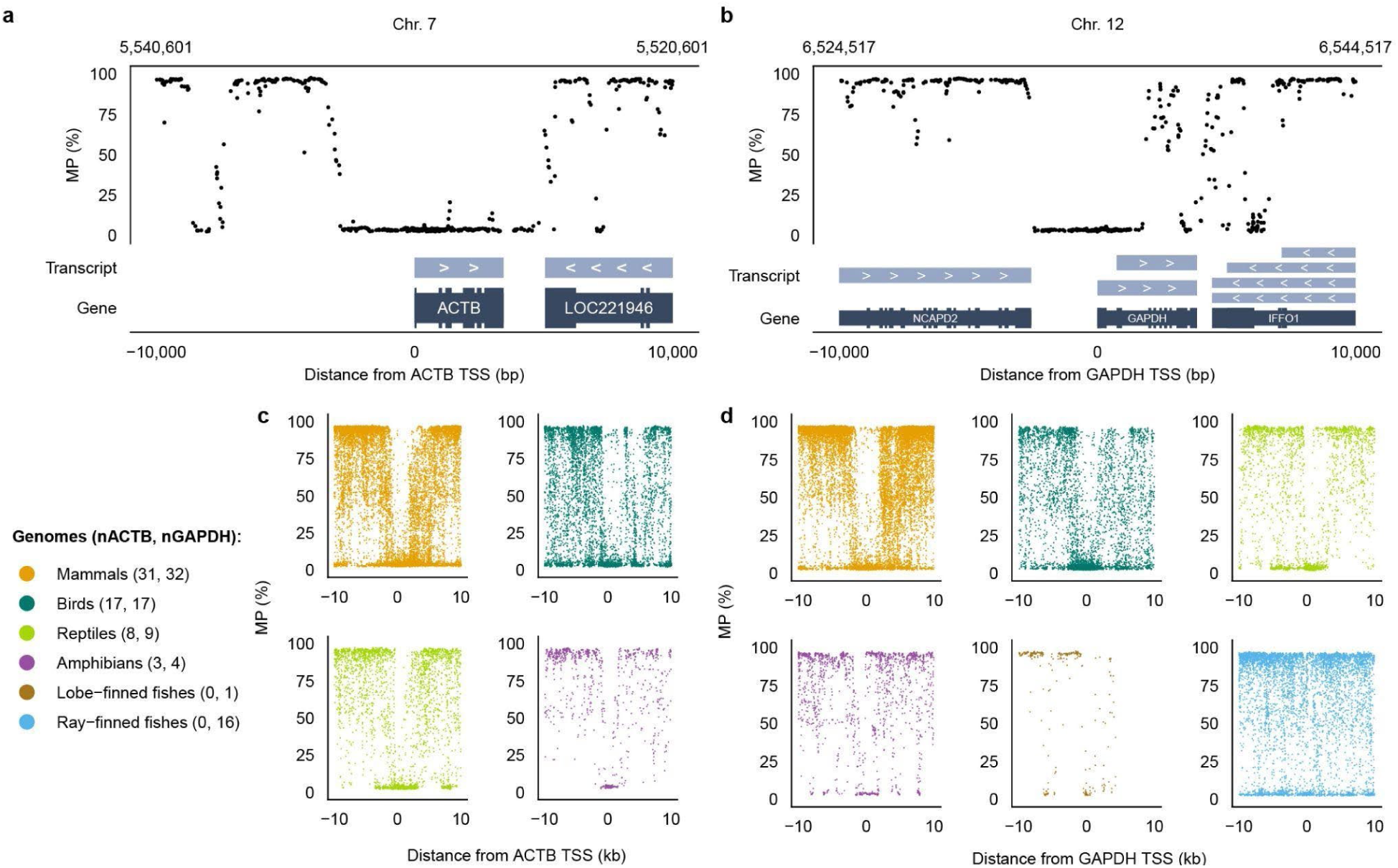
| Methylation profile at constitutively expressed genes. **a, b**, MP levels of CpGs on T2T-CHM13v2.0 genome with respect to physical distances from TSS for genes *ACTB* (**a**) and *GAPDH* (**b**) within ±10,000 bp interval of the site. **c, d,** MP levels of CpGs on genomes of 65 (*ACTB*) and 80 (*GAPDH*) VGP Phase I species with respect to physical distances from TSS for genes *ACTB* (n_Mammals_ = 31, n_Birds_ = 17, n_Reptiles_ = 8, n_Amphibians_ = 3) (**c**) and *GAPDH* (n_Mammals_ = 32, n_Birds_ = 17, n_Reptiles_ = 9, n_Amphibians_ = 4, n_Lobe-finned fishes_ = 1, n_Ray-finned fishes_ = 16) (**d**), separated into individual plots by class.

**Extended Data Fig. 8.**
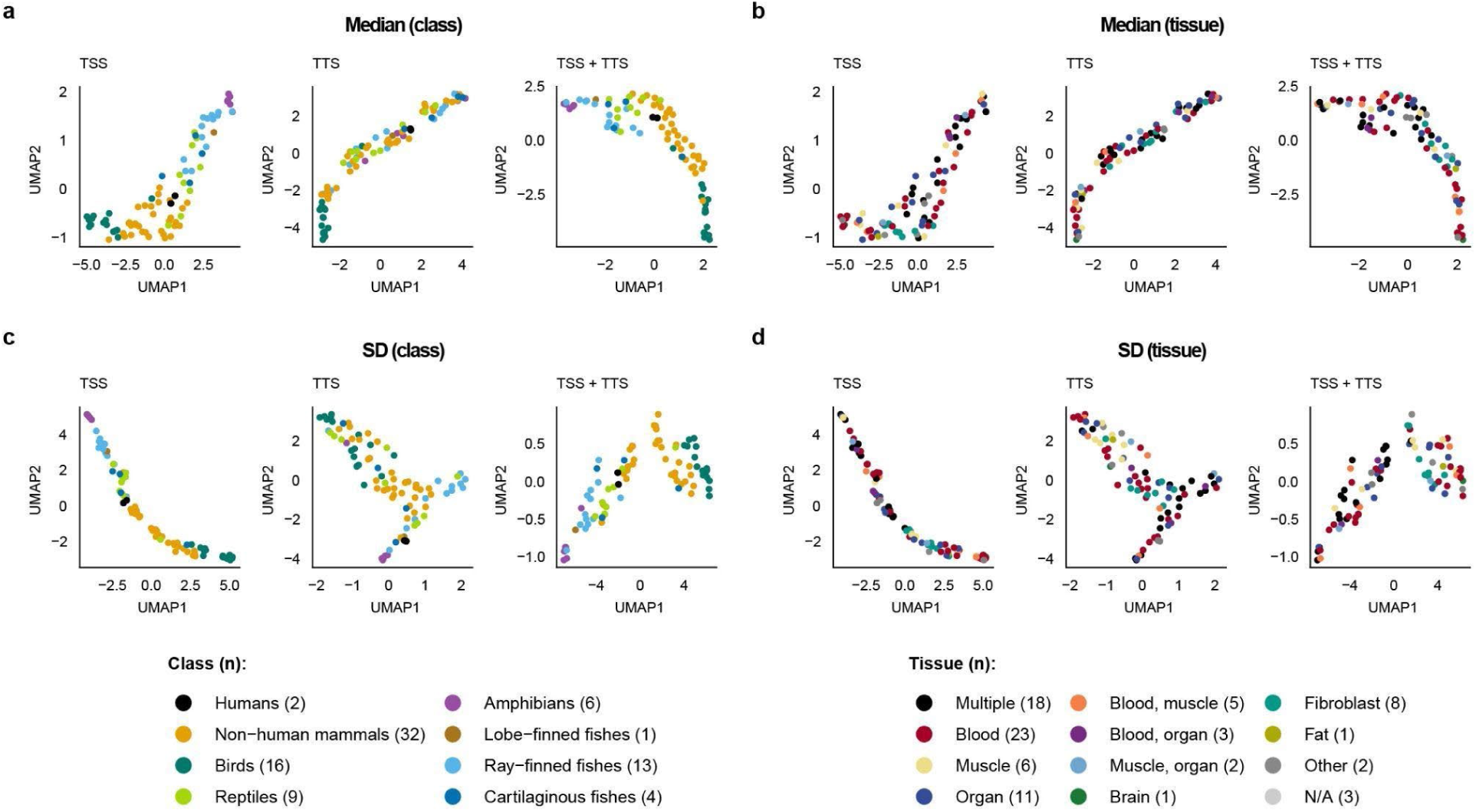
| Promoter-associated methylation clusters summarized in median and standard deviation clustered by lineage, more than by the tissue of origin. **a,b**, UMAP analysis of MP profiles summarized within ±10 kb of annotated transcription start sites (TSS), transcription termination sites (TTS), or both combined (TSS+TTS) across 90 VGP genomes. For each feature set, MP profiles were summarized by the median and standard deviation (left to right). Each point represents one genome, colored by taxonomic class using the same color codes as in Fig. 3. **c,d**. The same UMAP projections shown in **a,b** are colored by tissue of origin. Across feature sets and genomic contexts, samples separate primarily by lineage rather than tissue.

**Extended Data Fig. 9.**
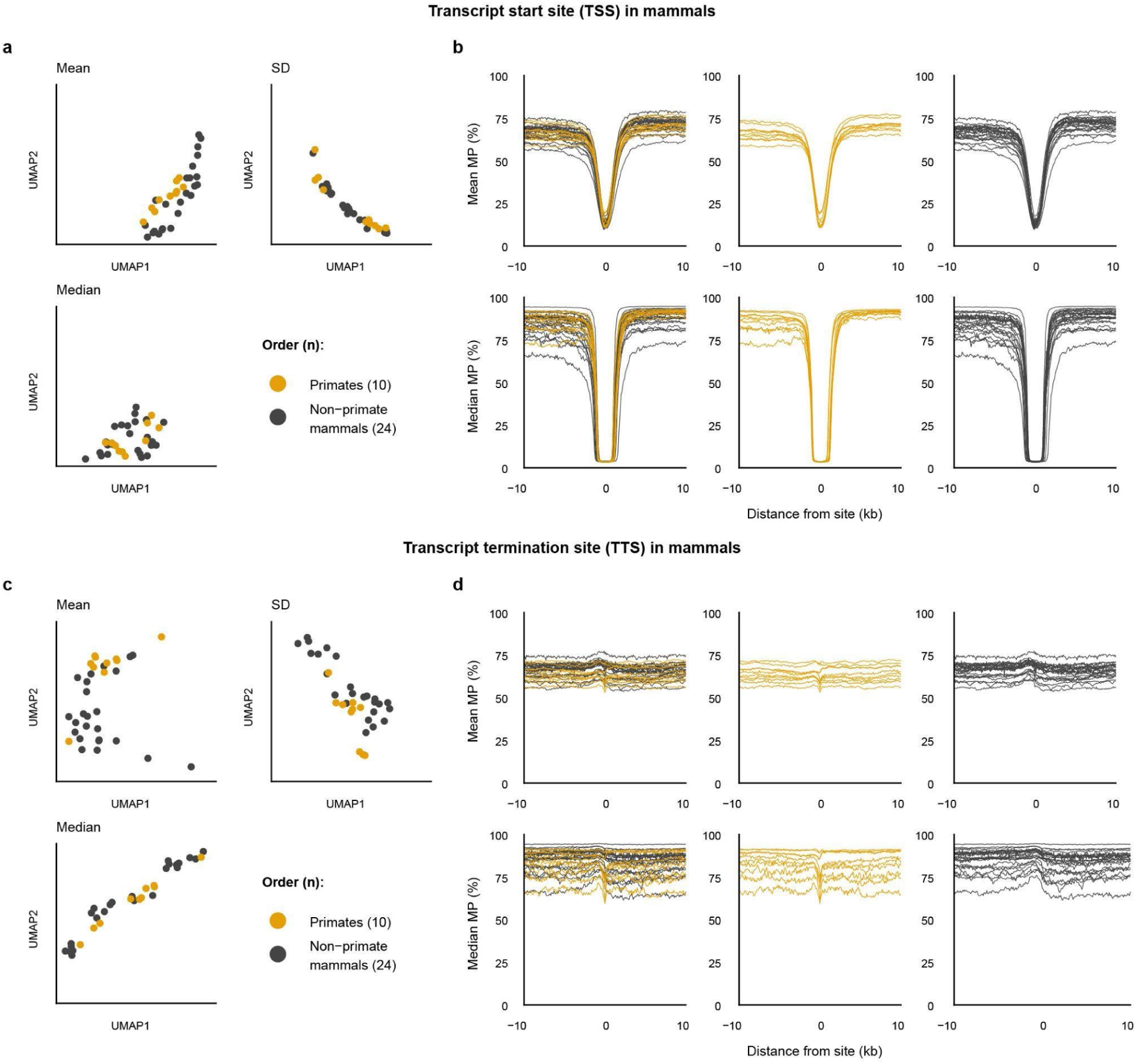
| Order-level classification of primates by mammal promoter methylation profiles. **a-d**, UMAP results based on mean, median, and standard deviation of MP profile at TSS (**a**) and TTS (**c**), and epigenetic discriminability of the primate order from the mammal class displayed by line plots of mean and median MP levels at TSS (**b**) and TTS (**d**).

**Extended Data Fig. 10.**
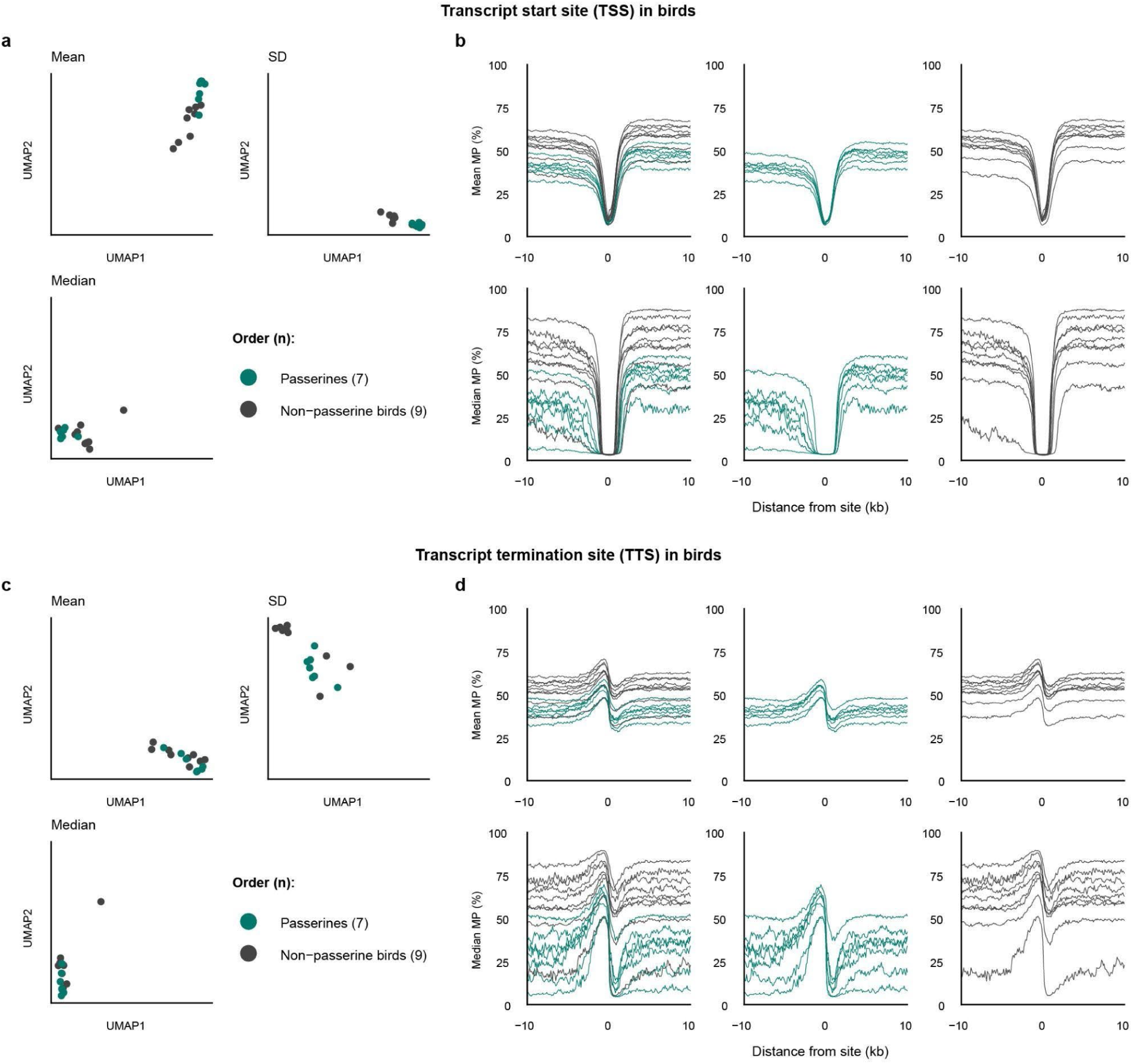
| Order-level classification of passerines by bird promoter methylation profiles. Identical format as **Extended Data** Fig. 9 for the passerine order from the bird class.

## Data availability

The RefSeq genomes used in this study are available in NCBI under the accession numbers listed in **Supplementary Table 1**. For methylation profiles, HiFi sequencing reads of human HG002 are available on the Data Explorer page of the Human Pangenome Resource Consortium (https://humanpangenome.org/).

HiFi sequencing reads of all other 87 assemblies used in this study are available in GenomeArk (http://genomeark.org) under assembly identifiers specified in **Supplementary Table 1**.

## Code availability

All codes were made available on GitHub (https://github.com/yh1126611/promoter_methylation_calculations).

## Acknowledgement

We would like to thank all members of the Vertebrate Genomes Project and the NCBI RefSeq eukaryote gene annotation team. We would like to thank Dr. Davenport for his insightful comments on the visualization items of this work. This work was supported by the National Research Foundation (NRF) of Korea grant funded by the Korea government (MSIT) (No. RS-2025-00513118 for Y.H.L. and H.K.), Republic of Korea. This work was financially supported by the NIH T-R01 (R01DC018691 for C.L., E.D.J.) and the Howard Hughes Medical Institute (E.D.J.), USA.

## Author information

These authors contributed equally: Young Ho Lee, Chul Lee

## Contributions

Y.H.L. and C.L. designed the study, collected data sets, conducted all analyses, and wrote the draft paper. All authors reviewed the draft. E.D.J. and H.K. secured funding. C.L., E.D.J., and H.K. co-supervised the study.

## Corresponding authors

Correspondence to Erich D. Jarvis and Heebal Kim

