## Supplementary material for "Epigenomic methylome landscape of promoters in vertebrate genomes": Online methods

### Data Download

*Homo sapiens* (human) sequencing data for sample HG002 were downloaded from Human Pangenome Reference Consortium[1] Release II Data Explorer (https://humanpangenome.org/). Human genomes hs1 (T2T-CHM13v2.0) and hg38 (GRCh38.p14) were downloaded from NCBI under accessions GCF_009914755.1 and GCF_000001405.40, respectively. The *Taeniopygia guttata* (zebra finch) genome for the previous reference assembly bTaeGut1.4.pri was downloaded from NCBI under accession GCF_003957565.2. Sequencing raw read data of all other VGP Phase I species in the dataset of this study (n=579 total) were downloaded from the repository of GenomeArk (http://genomeark.org) under assembly identifiers specified in **Supplementary Table 1**. All downloaded sequencing data were PacBio raw reads. We selected those that had BAM-formatted reads generated by CCS technology with the 5mC option, thus containing kinetics tags (forward interpulse duration, "fi"; forward pulse width, "fp"; reverse interpulse duration, "ri"; reverse pulse width, "rp"). We used 88 genomes that both meet this criterion and have primary assembly annotations from RefSeq. The reference genome assembly and the accompanying GTF-format RefSeq annotation for each species in our dataset were downloaded from NCBI under the accession numbers specified in **Supplementary Table 1**. All metadata regarding samples used in downstream analyses, such as tissue types, were extracted from respective NCBI genome pages. List of promoter (n=40,891) and enhancer (n=961,227) candidates on hg38 provided by the ENCODE database[2] was downloaded via the ENCODE Registry of cCREs V3, Search Candidate cis-Regulatory Elements (SCREEN) (https://screen.encodeproject.org/).

**Methylome assembly**

To construct the human methylome, we benchmarked a previously existing workflow[3] using updated versions of its software and algorithms. HG002 sequencing data was filtered for only HiFi (QV≥20, 0.99) reads by "extracthifi" tool of PacBio BAM toolkit (pbtk) v3.5.0 (https://github.com/PacificBiosciences/pbtk). 5mC methylation states were called from HiFi reads using pbjasmine v2.7.99 (https://github.com/PacificBiosciences/jasmine). Methylation-called reads were aligned to the hs1 genome using pbmm2 v26.1.0 (https://github.com/PacificBiosciences/pbmm2/), a PacBio wrapper for minimap2[4]. Aligned reads were sorted, and an index was generated by "sort" and "index" functions of samtools v1.23[5], respectively. MP (Methylation probability, 0-100%) at each genomic CpG was calculated by methylation states of reads mapped to the site and adjusted by a machine-learning algorithm based on genome-wide MP distribution using pb-CpG-tools v3.0.0 (https://github.com/PacificBiosciences/pb-CpG-tools) (**Fig. 1a**). This procedure was repeated with all sequencing data of all species (n=88) and the associated genome (n=90) in our dataset to construct methylomes for a total of 90 genomes.

Human genes were located on hs1 based on start and end coordinates of gene entries of its RefSeq annotation (GCF_009914755.1). Human centromere satellites were located using centromeric satellite annotation data[6] available at the UCSC Genome Browser. Hypo- and hypermethylated sites were defined as CpGs with MP < 10% and > 90%, respectively, as identified from our methylome assembly results. Four GC-comprised dinucleotides were located by a custom script "findseq.py" (https://github.com/yh1126611/promoter_methylation_calculations/) in Python 3.12.8. Coordinates of all located elements were converted to a format compatible with the downstream visualization tool by a custom script "bed_2_tbtools.py" (https://github.com/yh1126611/promoter_methylation_calculations/) in Python. Each element was visualized as a circos plot track using TBtools-II v2.420[7] software. (**Fig. 1h**). The visualization procedure was repeated with the T2T genome of zebra finch (bTaeGut7.mat) and its associated data (**Extended Data Fig. 1**).

Additionally, all GC content (**Fig. 1b**) as well as combined (**Fig. 1c**) and individual (**Fig. 1d**) frequencies of each of GC-comprised dinucleotides were calculated by a custom script "findseq.py" (https://github.com/yh1126611/promoter_methylation_calculations/) in Python3. All above observed genome-wide frequencies were compared with expected frequencies - 50% for GC contents [G, C] in [A, T, G, C]; 37.5% for GC-comprised dimers; 9.375% for CpG - and visualized as barplots using custom script "intro.R" (https://github.com/yh1126611/promoter_methylation_calculations/) employing packages scales 1.4.0[8] and ggplot2 3.5.2[9] in RStudio version: 2026.01.0+397[10] powered by R-3.6.0+[11]. Given that a base X in a genome has two juxtaposed dinucleotides (WXY), the expected probability of a base X constituting a GC-comprised dinucleotide was given by the sum of the probabilities of only WX being a GC-dimer (12.5%), only XY being a GC-dimer (12.5%), and all XYZ being a GC-dimer (12.5%), which is 37.5%. Likewise, the expected frequency of each GC-dimer was calculated as 1/4 of the expected frequency of all GC-dimers (9.375%). MP (0-100%) of genome-wide and chromosome-specific CpGs on hs1 were extracted from the methylome assembly result and visualized as a frequency histogram (**Fig. 1e**) and barplot indicating frequency (**Fig. 1f**) or chromosomal proportion (**Fig. 1g**) by height and MP by shade using a custom script "intro.R" (https://github.com/yh1126611/promoter_methylation_calculations) employing packages scales and ggplot2 in R.

**Methylation profiling at regulatory elements and various estimated promoter sites**

CpGs in ±10,000 bp vicinity of centers of all RefSeq-entered promoters (n=374), enhancers (n=107,386), silencers (n=34,232), and ENCODE promoter (n=40,891) and enhancer (n=961,227) candidates on hg38 were scanned for by "intersect" function of BEDtools v2.31.1[12,13], and their MP information was extracted from the human methylome previously assembled using hg38 as reference. Only the elements with one or more CpG in ±10,000 bp vicinity from the center (n = 373, RefSeq promoter; 107,163, RefSeq enhancer; 34,189 RefSeq silencer; 40,852 ENCODE promoter; 960,756, ENCODE enhancer) were retained. For each element, mean and median MP of all nearby CpGs ($n_{CpG}$ = 128,268, RefSeq promoter; 33,968,962, RefSeq enhancer; 12,378,864, RefSeq silencer; 14,648,009, ENCODE promoter; 256,216,476, ENCODE enhancer) at every base-unit distance was calculated for the interval ±10,000 bp. The resulting MP profiles were visualized as line plots showing distances from centers against the mean and median MPs of CpGs using a custom script, "plot_profile.R" (https://github.com/yh1126611/promoter_methylation_calculations), which employs the data package.table 1.17.8[14], dplyr 1.1.4[15], and ggplot2 in R (**Fig. 2a,b**).

To profile methylation properties at estimated promoter and transcription termination sites, we identified various start and end sites for transcription on hg38 based on gene (n=67,512) and transcript (n=202,461) entries using appropriate RefSeq annotations. Start sites (GSS, DGSS, ATSS, TSS) were each defined as 5'-termini of all genes, 5'-termini of genes not coinciding (discordant) with any transcript 5'-termini, 5'-termini of transcripts not coinciding with any gene 5'-termini, and 5'-termini of all transcripts, respectively. Similarly, end sites (GTS, DGTS, ATSS, TSS) were each defined as 3'-termini of all genes, 3'-termini of genes not coinciding with any transcript 3'-termini, 3'-termini of transcripts not coinciding (discordant) with any gene 3'-termini, and 3'-termini of all transcripts, respectively. Only the start and end sites with one or more CpG in ±10,000 bp vicinity (n = 32,562, GSS; 9,208, DGSS; 15,920, ATSS; 39,274, TSS; 32,522, GTS; 9,199, DGTS; 11,396, ATTS; 34,719, TTS) were retained for further analysis. Following the aforementioned visualization procedure, MP profiles shaped by CpGs ($n_{CpG}$ = 8,389,252, GSS; 1,822,736, DGSS; 4,603,235, ATSS; 11,170,600, TSS; 7,728,360, GTS; 2,444, DGTS; 2,718,948,

ATTS; 8,628,676, TTS) for all start and end sites of hg38 were generated as line plots (**Fig. 2c**, **Extended Data Fig. 2**).

The promoter and transcription termination site estimation was repeated on the human hs1 genome as well as the zebra finch genomes bTaeGut1.4.pri and bTaeGut7.mat to locate, profile, and visualize methylation of GSS, DGSS, ATSS, TSS, GTS, DGTS, ATTS, and TTS of each genome (**Extended Data Fig. 3,4,5**), with the number of sites and CpGs accounted in the calculations summarized in **Supplementary Table 2**.

The TSSs of all other genomes in our dataset (**Supplementary Table 1**) were located based on appropriate RefSeq annotations for each genome, and TSS methylation was profiled using the same procedure as for hg38, hs1, bTaeGut1.4.pri, and bTaeGut7.mat, using previously assembled methylomes for each species. To correct for interspecies differences in absolute MP levels, we applied a min-max normalization to the profile of each species, computing normalized MP for CpG $i$ ($MP_i^{norm}$) as:

$$MP_i^{norm} = \frac{MP_i - MP_{min}}{MP_{max} - MP_{min}} \times 100$$

with $MP_{min}$ and $MP_{max}$ representing the minimum and maximum MP for the corresponding species, respectively. Normalization did not alter the data trend, as all species exhibited $MP_{min}$ below 2% and $MP_{max}$ greater than or equal to 98% before normalization. Through visual inspection of profiles, six genomes manually judged to be exhibiting aberrant or artifactual MP patterns such as constant levels or mechanistic oscillation even after normalization - *Apodemus sylvaticus* (wood mouse), *Loxodonta africana* (African elephant), *Thunnus albacares* (yellowfin tuna), *Carassius carassius* (crucian carp), *Platichthys flesus* (European flounder), *Accipiter gentilis* (Northern goshawk), and zebra finch (bTaeGut7.mat genome only) - were excluded (n = 83; 36, Mammal; 18, Bird; 9, Reptile; 6, Amphibian; 1, Lobe-finned fish; 13, Ray-finned fish; 4, Cartilaginous fish).

**Significance calculation**

Given the non-normal nature of genomic MP distribution (**Fig. 1e**), a test for total variation distance ($d_{TV}$) was employed instead of the *t*-test for the deviation of MP at a certain site from the genome-wide distribution. Traditionally, $d_{TV}$ is defined as

$$d_{TV}(P, Q) = \frac{1}{2}\int |P(x) - Q(x)| dx$$ [16,17]

for two distributions $P$ and $Q$. For every $d_{TV}$ calculation, observed $d_{TV}(d_{TV,Obs.})$ between genome-wide CpGs ($P$) and CpGs at a given distance from the site ($Q$) was first approximated as

$$d_{TV,Obs.}(P, Q) = \frac{1}{2}\sum_{i=1}^{K} |\widehat{p_i} - \widehat{q_i}|$$

where $K$ is the number of bins for every base-unit distance within 10,000 bp from each site of interest. We set $K$ = 100 and divided all CpGs into 100 bins by MP. Then, expected $d_{TV}$ ($d_{TV,Exp.}$) values were estimated via 1,000 random permutations from resampling subsets of sizes equal to $Q$ from $P$, and their difference from $d_{TV,obs.}$ was used to compute empirical two-tailed p-values under the null hypothesis ($H_0$) that $P$ and $Q$ derive from identical distributions. $d_{TV}$ calculation was conducted on every site within ±10,000 bp from regulatory centers of hg38 (**Fig. 2a,b**) as well as transcription-related start and end sites of hg38 (**Fig. 2c, Extended Data Fig. 2**), hs1 (**Extended Data Fig. 3**), bTaeGut1.4.pri (**Extended Data Fig. 4**) and bTaeGut7.mat (**Extended Data Fig. 5**). Additionally, dTV calculation was conducted on all 90 genomes in our dataset (**Supplementary Table 1**) and class-pooled data consisting of and each class-pooled genome of mammal (**Fig. 6a**), bird (**Fig. 6b**), reptile (**Fig. 6c**), amphibian (**Fig. 6d**), lobe-finned fish (**Fig. 6e**), ray-finned fish (**Fig. 6f**) and cartilaginous fish (**Fig. 6g**) classes to compare distributions between 5'- and 3'-side of TSS to confirm statistical significance of epigenetic asymmetry suggesting a core promoter with smaller $K$ (30) to account for smaller $P$ and $Q$ sizes (number of CpGs). All $d_{TV}$ calculations were implemented using a custom script "compute_tvd.R" (https://github.com/yh1126611/promoter_methylation_calculations) employing package data.table and visualized as heatmaps indicating p-value at each distance through a custom script “visualize_tvd.R” (https://github.com/yh1126611/promoter_methylation_calculations) employing package ggplot2 in R.

**Methylation profiling at individual genes**

Coordinates of two housekeeping genes, *ACTB* and *GAPDH,* on hs1, their transcripts, and all neighboring (Distance ≤10,000 bp) genes and transcripts were identified from the RefSeq annotation. MP calculations of all CpGs within ±10,000 bp (n = 618, *ACTB*; 534, *GAPDH*) were extracted from our previously assembled human methylome. The CpGs were plotted as scatterplots with MP on the y-axis and distance from the GSS on the x-axis, using a custom script “scatterplot_gene.R” (https://github.com/yh1126611/promoter_methylation_calculations) employing the package ggplot2 in R (**Extended Data Fig. 7a,b**).

The above procedure was repeated on all species in our dataset that contained orthologs of *ACTB* (n=65) and *GAPDH* (n=80) (**Supplementary Table 1**). The data were classified by taxonomic class for each gene ($n_{ACTB}$ = 31, Mammal; 17, Bird; 8, Reptile; 3, Amphibian; $n_{GAPDH}$ = 32, Mammal; 17, Bird; 9, Reptile; 4, Amphibian; 1, Lobe-finned fish; 16, Ray-finned fish) for the scatterplots (**Extended Data Fig. 7c,d**).

**Core promoter breadth estimation**

After significance calculation of MP difference between 5’- and 3’-sides of TSS, core promoter breadth for each genome or class was defined as the furthest distance from TSS where all 10 bp-windows within the distance display 5’<3’ MP differences with significance (p<5%) confirmed by $d_{TV}$. This approach was applied to assemblies of all 82 species. Additionally, after pooling CpGs of all member species of each vertebrate class, excluding the hg1 and bTaeGut7.mat genomes to prevent repetitive incorporation of the same species, we calculated the average core promoter size for each vertebrate class. Core promoter sizes of genomes displaying 3'<5' MP differences from the window closest to TSS (0-10 bp) were labelled as undetectable ($n_{Undetectable}$ = 5, Bird; 2, Reptile; 1, Fish; 1, Cartilaginous fish). Core promoter sizes of genomes displaying significant 5'<3' MP differences in all windows inside the ±2,000 bp interval range from TSS were labelled as out of bounds of our calculation ($n_{OB}$ = 2, Bird; 1, Reptile) (**Supplementary Table 1**).

**Broader promoter breadth estimation**

Promoter start and end positions relative to TSS were estimated by first scanning median MP values at 10,000 bp upstream and downstream of all genomic TSS. Then, the difference between the mean of every median MP values at base-unit distances on promoter and non-promoter region within the ±10,000 bp window was calculated for all possible promoter delineations formed by combinations of promoter start position $i$ and end position $j$ where $i \geq -10{,}000$, $j \leq 10{,}000$, and $i < j$ was calculated by a custom script "promoter_delineation_calculator.py" (https://github.com/yh1126611/promoter_methylation_calculations) in Python. The delineation site combination with the largest mean difference between the median MP of the promoter and non-promoter regions was defined as promoter start and end sites. Median was used instead of mean for this calculation because the median profile showed a starker difference between baseline levels and promoter-associated drop. Only the promoter delineations encompassing TSS (position=0) and the core promoter (position = TSS - estimated core promoter size) were considered. This approach was applied to estimate promoter start and end sites and promoter sizes (end site - start site). Additionally, after pooling CpGs of all member species of each vertebrate class, excluding the hg1 and bTaeGut7.mat genomes to prevent repetitive incorporation of the same species, we calculated the average promoter size for each vertebrate class. All estimated core promoter and broader promoter breadths were visualized by a custom script "plot_profile_promoter.R" (https://github.com/yh1126611/promoter_methylation_calculations) employing ggplot2 package in R (**Fig. 5d-g, Fig. 6**).

The relationships between estimated promoter, core promoter and genome sizes of every species excluding species with undetectably small or over 10,000 bp promoter sizes, out of 2,000 bp range core promoter sizes or predefined outliers based on MP profiles ($n_{Out, P}$ = 20) (n=70) (**Supplementary Table 1**) were visualized as three scatterplots each with genome size/promoter size (**Figure 5h**), genome size/core promoter size (**Figure 5i**), and promoter size/core promoter size (**Figure 5j**) on X and Y-axes, respectively, by a custom script "plot_comparison.R" (https://github.com/yh1126611/promoter_methylation_calculations) employing ggplot2 package in R. Genome sizes were defined as the sum of the lengths of all placed chromosomes on the RefSeq genome (**Supplementary Table 1**).

**UMAP of the vertebrate genome dataset based on promoter methylation**

Uniform manifold approximation and projection (UMAP) analysis was conducted on all genomes of our dataset (n=83). The mean, median, and standard deviation (SD) of MP at each base-unit distance from TSS inside a ±10,000 bp interval ($n_{MP.}$ = 20,001 for each genome) were used as input data to the R function umap from the package uwot 0.2.4[18] with n_neighbors=15 and n_components=2 to generate component values. UMAP results for the top two components (UMAP1 and UMAP2) were visualized using a custom script, "plot_umap.R" (https://github.com/yh1126611/promoter_methylation_calculations), which employed the ggplot2 package in R (**Fig. 4, Extended Data Fig. 8**).
